# Diet Post-translationally Modifies The Gut Microbial Proteome To Modulate Renal Function

**DOI:** 10.1101/2020.02.25.964957

**Authors:** Lior Lobel, Y. Grace Cao, Jonathan N. Glickman, Wendy S. Garrett

## Abstract

We identify a novel mechanism linking diet, gut microbial metabolism, and renal function. We found that a sulfur amino acid-based dietary intervention post-translationally modifies a microbial enzyme, blunting its uremic toxin-producing activity and alleviating chronic kidney disease (CKD) in a preclinical model. We also define a heretofore unknown role for the post-translational modification S-sulfhydration within the gut microbiome. This study provides a framework for understanding how diet can tune microbiota function via protein post-translational modification without altering microbial community composition to support healthy host physiology beyond the gut and specifically how a dietary modification can inhibit tryptophanase activity to ameliorate CKD progression.

**One Sentence Summary:** We found that diet post-translationally modifies the gut microbiota proteome to modulate kidney function.

## Main Text

Chronic kidney disease (CKD) affects nearly 850 million people worldwide (*1*). Although dietary modification is a cornerstone of CKD treatment, the mechanistic roles of diet-microbiota interactions in CKD pathogenesis and treatment have been under-explored. While many diet-microbiome studies have focused on the effects of dietary fiber, fat and carbohydrates (*2*), less is known about the specific effects of dietary protein and amino acids, although 5-10% of dietary amino acids reach the colon where most gut bacterial metabolism occurs (*3*). In humans, increasing dietary protein increases gut bacterial production of hydrogen sulfide (H_2_S), indole, and indoxyl sulfate (*4, 5*). Indole and indoxyl sulfate are uremic toxins; and H_2_S has diverse physiological functions, some of which are mediated by the post-translational modification S-sulfhydration (*6, 7*). While a vast number of studies have been performed in mammalian systems, the physiological roles of H_2_S in regulating gut bacterial function within a host are understudied. Additionally, whether there are *bona fide* opportunities to improve CKD by manipulating diet-microbiota interactions remain unclear.

Given the knowledge gaps around dietary protein, gut microbial metabolism and H_2_S, and to address the role of gut microbial metabolism and diet in renal function; we employed a mouse model of CKD that is driven by elevated adenine (*8*) along with a sulfur amino acid (Saa)-based diet perturbation. We formulated isocaloric diets to represent edge cases of mouse Saa consumption, i.e. diets with low versus high amounts of methionine and cysteine (see Table S1 for diet formulations) but with sufficient methionine to avoid methionine restriction (*9*). Conventionally-reared, specific pathogen-free (SPF) mice on a low Saa + adenine (Saa+Ade) diet had significantly increased serum creatinine levels compared to mice on high Saa+Ade (Fig. 1A), as well as more extensive and severe renal cortex histopathologic changes, including tubular dilatation and drop-out, tubulitis with peri-tubular fibrosis, and cortical crystal deposition (Fig. 1B-D). To determine the extent to which the Saa effects were dependent upon the gut microbiota, we fed the Saa+Ade diets to gnotobiotically-reared, germ-free (GF) mice. Serum creatinine and kidney damage were markedly reduced in the GF mice as compared to SPF mice on the low Saa+Ade diet, while there were similar phenotypes in the GF and SPF mice fed the high Saa+Ade diet (Fig. 1A-D). Overall, we found that a low Saa diet exacerbated the CKD phenotypes observed and the presence of a gut microbiota further magnified these effects.

**Fig 1.**
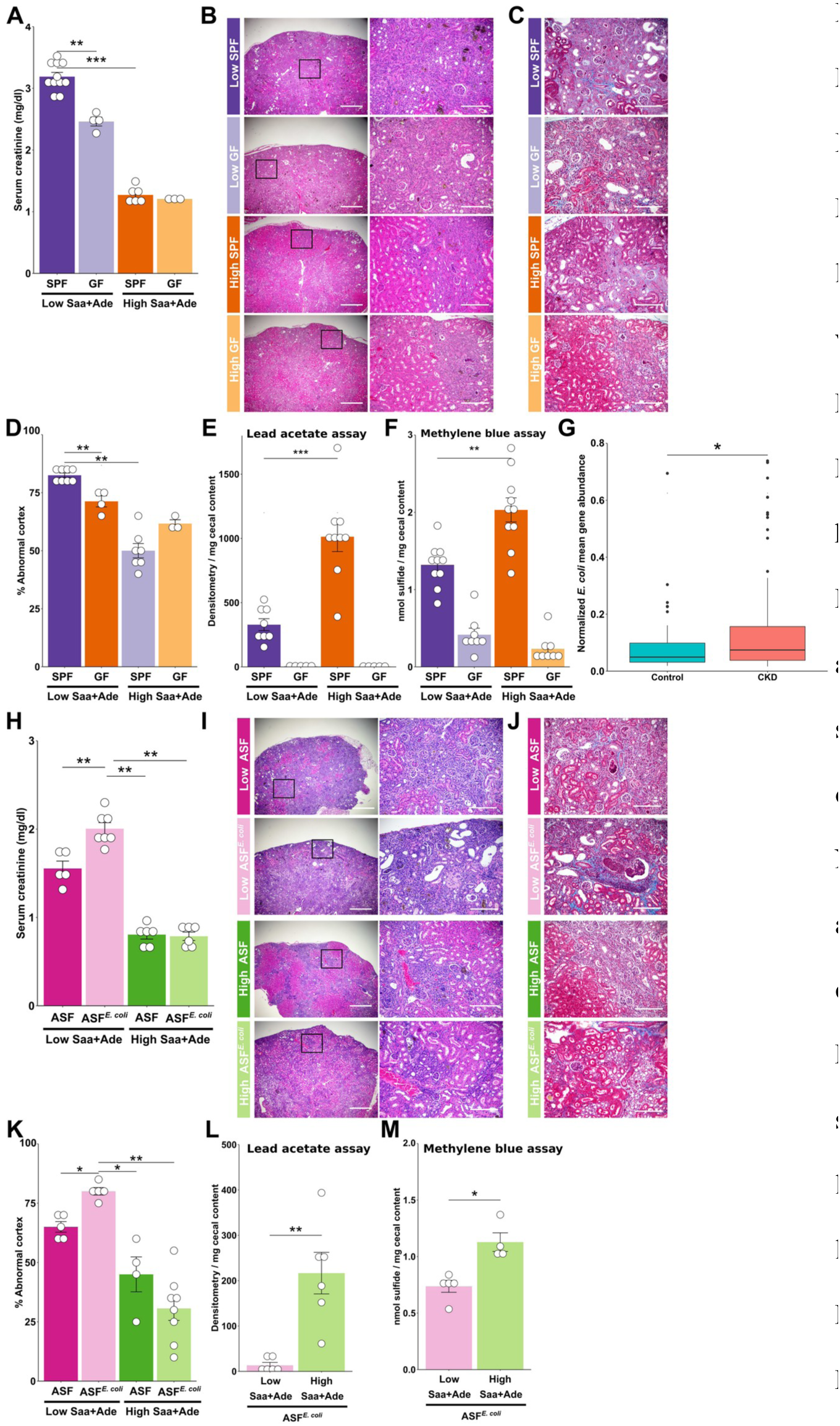
Dietary Saa and the Gut Microbiota Modulate Kidney Injury Severity in a Mouse CKD Model. A. Serum creatinine (Cre) levels of SPF and GF mice on low vs. high Saa+Ade diets. **B.** Representative H&E staining and **C.** Representative trichrome staining of kidneys from mice in A. **D.** Histology-based renal injury score. **E and F.** SPF and GF mice cecal sulfide levels detected by lead acetate or methylene blue assay. **G.** Normalized *E. coli* mean gene abundance in CKD patient samples compared to non-CKD controls, PTRI whole genome shotgun sequencing dataset. **H.** Serum Cre levels from ASF or ASF*^E. coli^* mice on low vs. high Saa+Ade diets. **I.** Representative H&E staining and **J**. Representative trichrome staining of kidneys from mice in H. **K.** Histology-based renal injury score. **L and M.** ASF and ASF*^E. coli^* cecal sulfide levels detected by lead acetate or methylene blue assay. Data represent 2 independent experiments for **L** and **M**, 3 for **A**, **D, H** and **K,** and 4 for **E** and **F.** Symbols represent individual mice. Bars represent mean ± SEM. * P value < 0.05, ** P value < 0.01, *** P value < 0.001. Two-way ANOVA with Tukey’s post-hoc test for **A, D**, **E, F, H** and **K**, and Mann-Whitney test for **L** and **M.**

A plausible link between dietary Saa and gut bacteria is microbial metabolism of cysteine to H_2_S. We measured cecal sulfide levels from GF and SPF mice fed low versus high Saa diets using both the lead acetate and methylene blue sulfide detection assays (*10*). SPF mice on the high Saa diet had higher cecal sulfide levels than those on the low Saa diet (Fig. 1E-F). GF mouse ceca had significantly less sulfide than SPF mice, regardless of Saa diet (Fig. 1E-F). We did not observe any significant differences in the taxonomic abundances of the gut microbiota members between SPF mice on the low vs high Saa diets using 16S rRNA gene amplicon surveys (Fig. S1), supporting that the differences in cecal sulfide in healthy mice may be mediated by altering microbial function, rather than changing microbiota community structure.

Given these findings and with the goal of more effectively modeling gut microbial activity shifts that could occur in CKD patients, we sought out publicly available CKD patient gut microbiota profiling studies to identify taxa enriched in CKD patients as compared to healthy individuals. We re-analyzed fecal 16S rRNA gene amplicon datasets from Xu *et al*. (*11*) and from Southern Medical University (NCBI accession PRJEB5761), a fecal PhyloChip study from Vaziri *et al*. (*12*), and a fecal whole genome shotgun sequencing dataset from Promegene Translational Research Institute (PTRI) (NCBI accession PRJNA449784). Enforcing stringent statistical cutoffs (LDA > 4 for LEfSe analyses and fold change > 2 for the PhyloChip analysis) revealed a clear and robust signal of *Enterobacteriaceae* enrichment in CKD patients (Fig. S2A-C). Although the 16S rRNA gene amplicon analyses did not afford species level *E. coli* identification, the PhyloChip analysis showed a significant increase in the combined mean abundance of seven *E. coli* strains measured in fecal samples of CKD patients with end-stage renal disease compared to control subjects (Fig. S2D). Further analysis of the PTRI whole genome shotgun sequencing dataset strengthened this finding, as we found a higher normalized *E. coli* mean gene abundance in CKD patient samples compared to non-CKD controls (Fig. 1G). Given these human CKD gut microbiota reanalysis findings and both the genetic tractability and relatively well-characterized proteome of *E. coli*, we focused on the effects of *E. coli* in the adenine-driven CKD model. As the mice we obtain from Jackson Laboratory do not harbor any *Enterobacteriaceae* members (Fig. S1) and to carry out a carefully controlled study of gut microbial activity in a reproducible model system we used gnotobiotically-reared mice colonized with the altered Schaedler Flora (ASF) to which we added *E. coli* K-12. The ASF is a simplified microbial community consisting of 8 bacterial species, none of which are related to *Enterobacteriaceae* (*13*). We employed ASF mice, rather than mono-colonized mice, because ASF mice are more physiologically similar to SPF mice (*13*). *E. coli* colonization was similar on low and high Saa and Saa+Ade diets (Fig. S3A-B), and we did not observe changes in the relative abundance of ASF members (Fig. S3C). On the low Saa+Ade diet, ASF mice colonized with *E. coli* (ASF*^E. coli^*) had higher serum creatinine and more extensive tubulitis, tubular atrophy and drop-out, peritubular fibrosis, and cortical crystals than ASF mice (Fig 1. H-K). In contrast, ASF*^E. coli^* and ASF mice on the high Saa+Ade diet had similar serum creatinine levels and milder renal parenchymal pathology compared with their littermates on the low Saa+Ade diet (Fig 1. H-K). As with SPF mice, we found higher cecal sulfide levels in ASF*^E. coli^* mice on the high versus low Saa+Ade diet (Fig 1. L-M). To determine if changes in renal function would occur in these models in the absence of the adenine insult, we examined creatinine levels in ASF*^E. coli^* mice on the low versus high Saa diet. Remarkably, the low Saa diet and *E. coli* were sufficient to increase serum creatinine levels in mice and no overt histologic abnormalities were present (Fig. S3D). Overall, these results support that *E. coli* interacts with dietary Saa to modulate kidney function.

Given our observations regarding cecal H_2_S in SPF and ASF*^E. coli^* mice on the Saa diets and the literature on how H_2_S can post-translationally modify mammalian proteins leading to a range of physiologic effects, we delved into examining the effects of H_2_S on *E. coli*. In lead acetate sulfide detection assays, *E. coli* produced sulfide from cysteine in a dose-dependent manner, grown aerobically or anaerobically, without any effects on growth (Fig. 2A and Fig. S4A-C). To serve as a control for how endogenous H_2_S production affects *E. coli* physiology, we generated an isogenic strain harboring a deletion of *decR*, which encodes a transcriptional activator of the cysteine de-sulfhydrase *yhaOM*, which drives cysteine-derived sulfide production in *E. coli* (*14*). *decR* deletion resulted in significant reduction of sulfide production, with no effect on growth kinetics (Fig. 2A-B and Fig. S4A-C). Sulfide exerts its effects through generation of polysulfides that modify cysteine residues, resulting in S-sulfhydration (*15*). To identify *E. coli* proteins that are S-sulfhydrated (R-S-S), we adapted a pull-down method that specifically enriches for S-sulfhydrated proteins (*16*). This technique leverages maleimide binding to free thiols, resulting in thioester bonds, and the ability of dithiothreitol (DTT) to break disulfide bonds but not thioester bonds (Fig. 2C) We observed a robust enrichment of S-sulfhydrated proteins in DTT-eluted samples using this method on WT *E. coli* lysates grown in media supplemented with cysteine (Fig. 2D). We validated the pull-down assay’s specificity and found that treating bacterial lysates with H_2_O_2_, and hence oxidizing free thiols, reduced the detection of S-sulfhydrated proteins (Fig. S4D). In contrast, treatment with sodium hydrosulfide (NaHS), a fast-reacting sulfide donor, induced higher S-sulfhydration levels in bacterial lysates (Fig. S4D). We detected a higher level of S-sulfhydration in *E. coli* lysates grown in media supplemented with cysteine compared to *E. coli* grown in LB alone (Fig S4E-F). In contrast, lysates of Δ*decR* bacteria, which produce less H_2_S, grown in cysteine-supplemented LB broth had lower S-sulfhydration than WT *E. coli* (Fig. 2E). Next, we sought to characterize the *E. coli* sulfhydrome using quantitative tandem mass tag (TMT) LC-MS^3^ analysis. This analysis revealed that most identified proteins were indeed S-sulfhydrated, as they were enriched in *E. coli* lysates that were eluted with DTT versus the same samples not treated with DTT (Fig. 2F). Furthermore, most detected S-sulfhydrated proteins were enriched in WT vs *ΔdecR E. coli*, as expected from the strains’ differential ability to produce sulfide from cysteine. Ranking of the S-sulfhydrated proteins by their q values (DTT versus non-DTT) revealed the top 10 most abundant S-sulfhydrated proteins (Fig. 2F and Fig. S4G). While most of these proteins are highly expressed during logarithmic bacterial growth, and are expected to be highly abundant, tryptophanase (TnaA) was over-represented. Overall, our quantitative proteomics analysis identified 212 proteins as S-sulfhydrated with high confidence (Table S2), and hyper-geometric distribution analysis revealed thirteen cellular pathways enriched with S-sulfhydrated proteins, several of which are related to protein translation (Fig. S4H).

**Fig 2.**
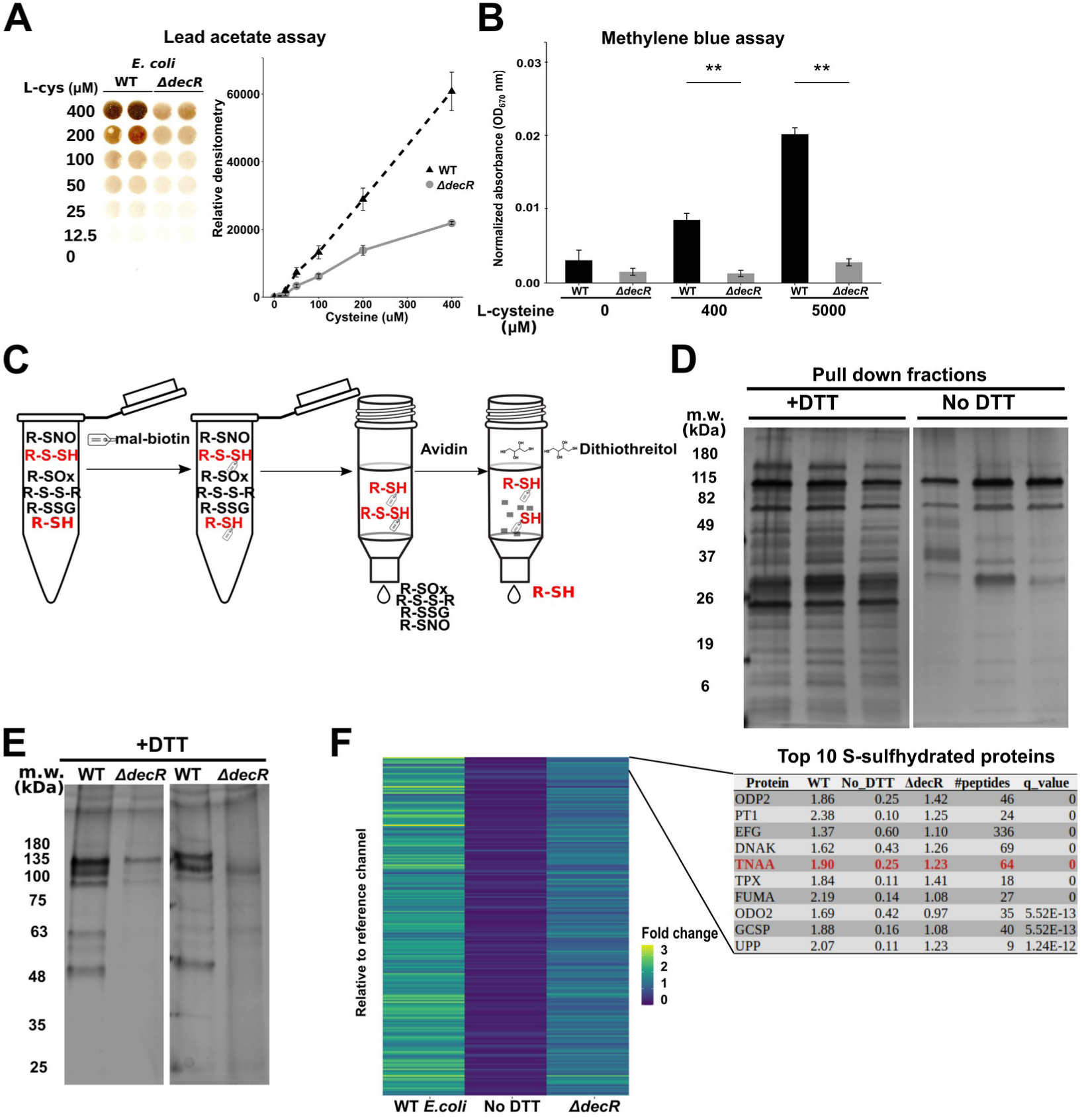
Characterization of *E. coli* S-Sulfhydrome Reveals that TnaA is a Highly S-Sulfhydrated Protein. A. *E. coli* sulfide production by lead acetate. **B.** *E. coli* sulfide production by methylene blue. **C.** Schematic of S-sulfhydrated protein pull-down method. **D.** Silver staining of *E. coli* lysates subjected to S-sulfhydration pull-down and eluted either with or without DTT. **E.** Silver staining of WT and Δ*decR E. coli* lysates subjected to S-sulfhydration pull-down. **F.** Heatmap of the relative quantity of the 212 S-sulfhydrated proteins by TMT LC-MS^3^ analysis from S-sulfhydration pull-down fractions from WT *E. coli* samples eluted with or without DTT and *ΔdecR* mutant samples eluted with DTT. Proteins ordered based on q-value score for enrichment in the DTT vs non-DTT eluted samples. Data represent 2 independent experiments for **E**, 3 for **D** and **F,** 4 for **A** and 6 for **B**. Bars represent mean ± SEM. ** P value < 0.01. Linear model test **A,** two-way Kruskal-Wallis test with Dunn’s post-hoc test **B** and two-way ANOVA with Tukey’s post-hoc test **F**.

To connect our S-sulfhydrome analysis to the phenotypes we observed in the CKD preclinical model, we focused on TnaA (Fig. 2F), a secreted enzyme that catalyzes the degradation of tryptophan to indole, pyruvate, and ammonia. Indoles are a class of bacterial-produced molecules that not only regulate bacterial physiology (*17*), but also participate in bacteria-host interactions (*18*). Indoles can be transported through the portal vein to the liver where they are oxidized, yielding the uremic toxin indoxyl sulfate (*19*). For these reasons, TnaA emerged as an attractive target for investigating host-microbe interactions in our CKD mouse model. We replaced the *E. coli* TnaA chromosomal copy with a cloned *tnaA-his* under its native promoter. We then validated our S-sulfhydrome results by analyzing TnaA S-sulfhydration in WT versus *ΔdecR E. coli* lysates using Western blot analysis and found reduced TnaA S-sulfhydration in *ΔdecR* lysates (Fig. 3A). *E. coli* lysates treated with H_2_O_2_ and NaHS showed reduced and increased TnaA S-sulfhydration, respectively (Fig. 3B). Since the S-sulfhydration pull-down method reduces the S-sulfhydrated cysteine residue (*i.e*. removes the S-sulfhydration), we could not pinpoint the exact cysteine residues being S-sulfhydrated, as TnaA has 7 cysteines. Therefore, we purified natively expressed TnaA-His from *E. coli* grown in LB supplemented with cysteine and performed LC-MS/MS analysis to detect and map the S-sulfhydration. We detected several TnaA-His peptides that had a +32 Da addition, matching the molecular weight of S-sulfhydration on a cysteine residue (Fig. S5). As oxidation of a cysteine residue to sulfinic acid (R-S-O_2_) results in same mass shift and given the potential for oxidation during our analysis, we could not rule out that such oxidation occurs. However, an S-sulfhydrated cysteine can be oxidized to sulfinic acid (R-S-S-O_2_) resulting in a +64 Da increase, a shift that results from oxidation of an S-sulfhydrated cysteine or a second S-sulfhydration (R-S-S-SH). We were able to detect a +64 Da shift in several cysteine residues of TnaA (Fig. S5 and Table S3). While we found evidence that 6 out of the 7 TnaA cysteine residues were S-sulfhydrated (C148, C281, C294, C298, C352 and C383), we cannot rule out that cysteine residue (C413) can also be S-sulfhydrated, as our coverage of TnaA (∼78%) did not include peptides with high confidence within this region. TnaA cysteine residues have been reported to be important for its enzymatic activity (*20*), as mutation of cysteine 298 results in inhibition of TnaA activity (*21*). To study the effect of S-sulfhydration on TnaA activity, we measured indole concentrations by both Kovac’s reagent and LC-MS/MS analysis of bacterial cultures. We found that supplementing LB broth with cysteine or NaHS reduced indole concentrations in the supernatants (Fig. 3C-D). Also, supporting sulfide’s role in TnaA inhibition, Δ*decR E. coli* had higher indole levels compared to WT *E. coli* when grown in LB supplemented with cysteine (Fig. S6A), and TnaA expression was similar under these conditions (Fig. S6B). To demonstrate that indole production was dependent on TnaA, we ablated TnaA activity by using an isogenic *tnaA739::kan* mutant (*tnaA mut*) and did not detect indole in the culture supernatant (Fig. S6C). To test that S-sulfhydration inhibits TnaA activity in a direct manner, we employed a reductionist approach using purified *E. coli* TnaA. We observed that incubation with disodium tetrasulfide (Na_2_S_4_), a poly-sulfide donor, led to TnaA S-sulfhydration (Fig. S6D) and reduced enzymatic activity by 60% *in vitro* (Fig. 3E). As an assay control, we added DTT, which should reduce S-sulfhydrated TnaA to its functional native form and observed TnaA activity more than triple (Fig. 3E). Collectively, these results support that S-sulfhydration of *E. coli* TnaA reduces its activity as measured by indole production from tryptophan, both *in vitro* and in bacterial cultures.

**Fig. 3.**
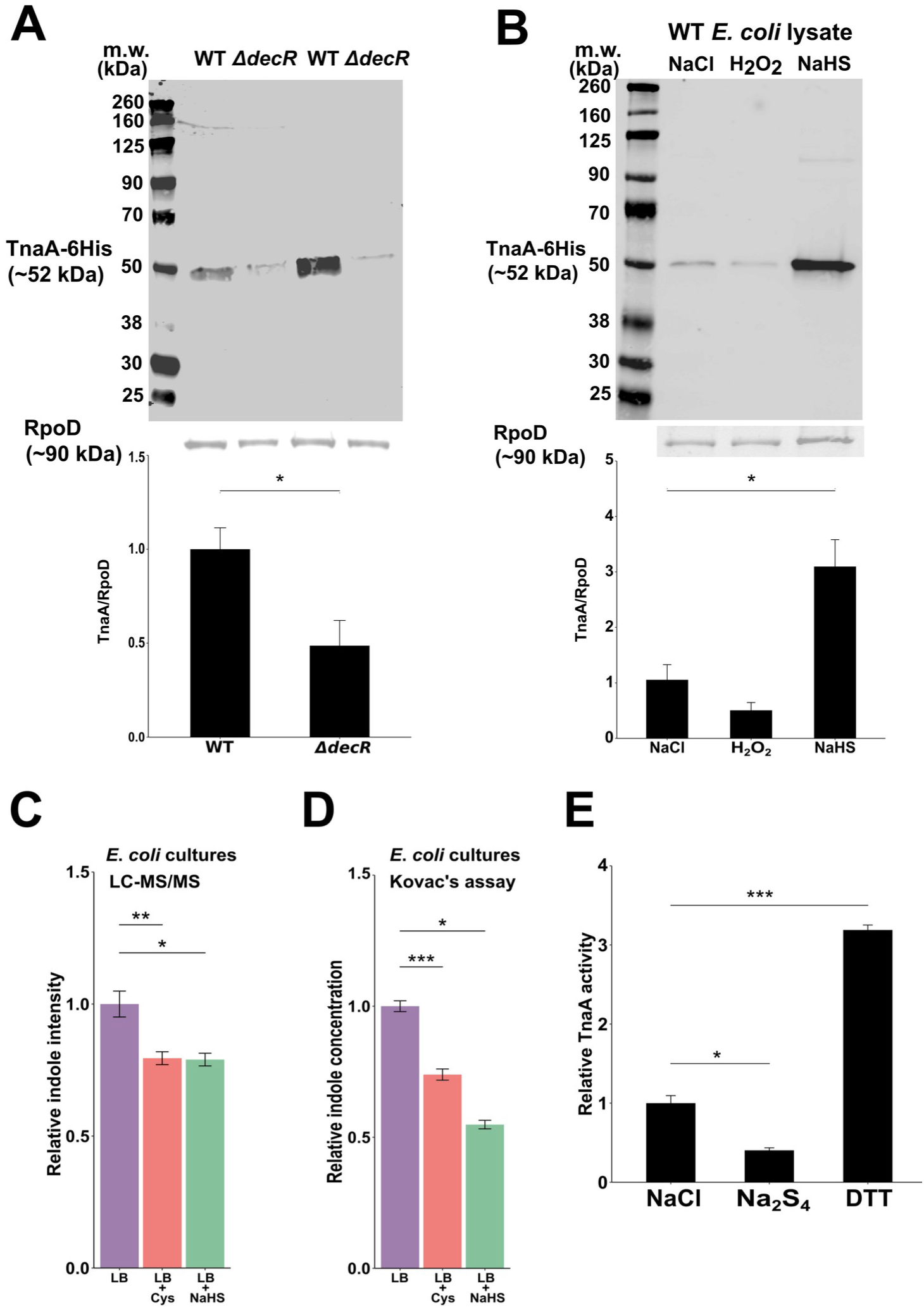
S-sulfhydration Inhibits the Indole-Producing Enzymatic Activity of *E. coli* TnaA. A. Representative western blot analysis of TnaA-His from WT and Δ*decR E. coli* lysates subjected to S-sulfhydration pull-down. Loading controls show RpoD in the flow-through. **B.** Same method as in **A** with *E. coli* lysates treated with NaCl, H2O2, or NaHS. **C.** LC-MS/MS analysis of indoles in WT *E. coli* cultures with cysteine or NaHS. **D-E.** Kovac’s assay for indole production in (**D**) WT *E. coli* cultures with cysteine or NaHS and (**E**) purified TnaA enzyme supplemented with NaCl, Na2S4, or DTT. Data represent 3 independent experiments for **A** and **E**, 4 for **B** and **D,** and 5 for **C**. Bars represent mean ± SEM. * P value < 0.05, ** P value < 0.01, *** P value < 0.001. Mann-Whitney test **A** and two-way ANOVA with Tukey’s post-hoc test **B** and **E,** and two-way Kruskal-Wallis test with Dunn’s post-hoc test **C** and **D**.

While we detected TnaA S-sulfhydration *in vitro* for both purified protein and TnaA from bacterial cell lysates and demonstrated that this modification inhibited its activity, we had yet to determine if this post-translational modification occurred within the gut in response to dietary Saa and resulted in measurable physiological consequences for the host. We began by providing ASF*^E.coli^* mice with the high and low Saa diets. While mice on the diets harbored similar levels of *E. coli* (Fig. S7A), we detected higher TnaA S-sulfhydration in the cecal contents of mice on the high Saa diet compared to those on the low (Fig. 4A). None of the 8 ASF bacterial genomes encode a *tnaA* gene, and we could not detect indoles in the ASF mouse cecal contents using LC-MS/MS, implying that *E. coli* is the sole producer of indoles in this model (Fig. S7B). Taking advantage of that distinction, we measured indole in the cecal contents of ASF*^E. coli^* mice on the two diets and found that mice on the high Saa diet had significantly lower indole levels demonstrating that high dietary Saa not only increased TnaA S-sulfhydration but also that this modification was sufficient to affect TnaA activity *in vivo* (Fig. 4B-C). To strengthen the links between diet, microbial metabolism, and kidney damage, we leveraged the CKD preclinical model using the low Saa+Ade diet, in which we observed the most renal injury and the gnotobiotic ASF mice we used previously (Fig. 1). We colonized these mice with either WT *E. coli (*ASF*^E.coli^*), or with one of two isogenic mutants, *tnaA mut* or Δ*decR (*ASF *^tnaA mut^* and ASF^Δ*decR*^, respectively). Colonization of the three different *E. coli* strains was similar (Fig. S7C). Unlike in ASF*^E.coli^* mice, no indoxyl sulfate was detected in the serum of ASF*^tnaA mut^* mice as there was no tryptophanase present within the gut microbiota. As *E. coli* Δ*decR* is deficient in producing sulfide from cysteine, TnaA remains less S-sulfhydrated/more highly active (Fig. 3A and S6A). Consistent with that observation, serum indoxyl sulfate from ASF^Δ*decR*^ mice was increased relative to the sera levels observed from ASF*^E. coli^* mice (Fig. 4D). Mice colonized with WT *E. coli* had higher serum creatinine levels compared to mice colonized with the *tnaA mut* strain (Fig. 4E). Concomitant with the serum indoxyl sulfate levels, mice colonized with the Δ*decR* strain had the highest serum creatinine levels (Fig. 4E). Histological findings of more severe tubulointerstitial damage, fibrosis, and cortical crystal deposition and more extensive parenchymal involvement mirrored the trends observed for indoxyl sulfate and creatinine for the *E. coli* Δ*decR* vs WT *E. coli* (Fig. 4F-G). We also examined these mice on the high Saa+Ade diet. Consistent with prior observations (Fig. 1), ASF*^E.coli^* mice on this diet demonstrated milder phenotypes, although ASF^Δ*decR*^ mice had slightly increased serum creatinine compared to the parental and *tnaA mut* strains (Fig. S7D-H). Collectively, these data support that a dietary component can generate a post-translational modification of a gut microbial protein that affects extra-intestinal host function and furnish mechanistic insight into how host-diet-microbiota interactions can contribute to prevalent disease states such as CKD.

**Fig. 4.**
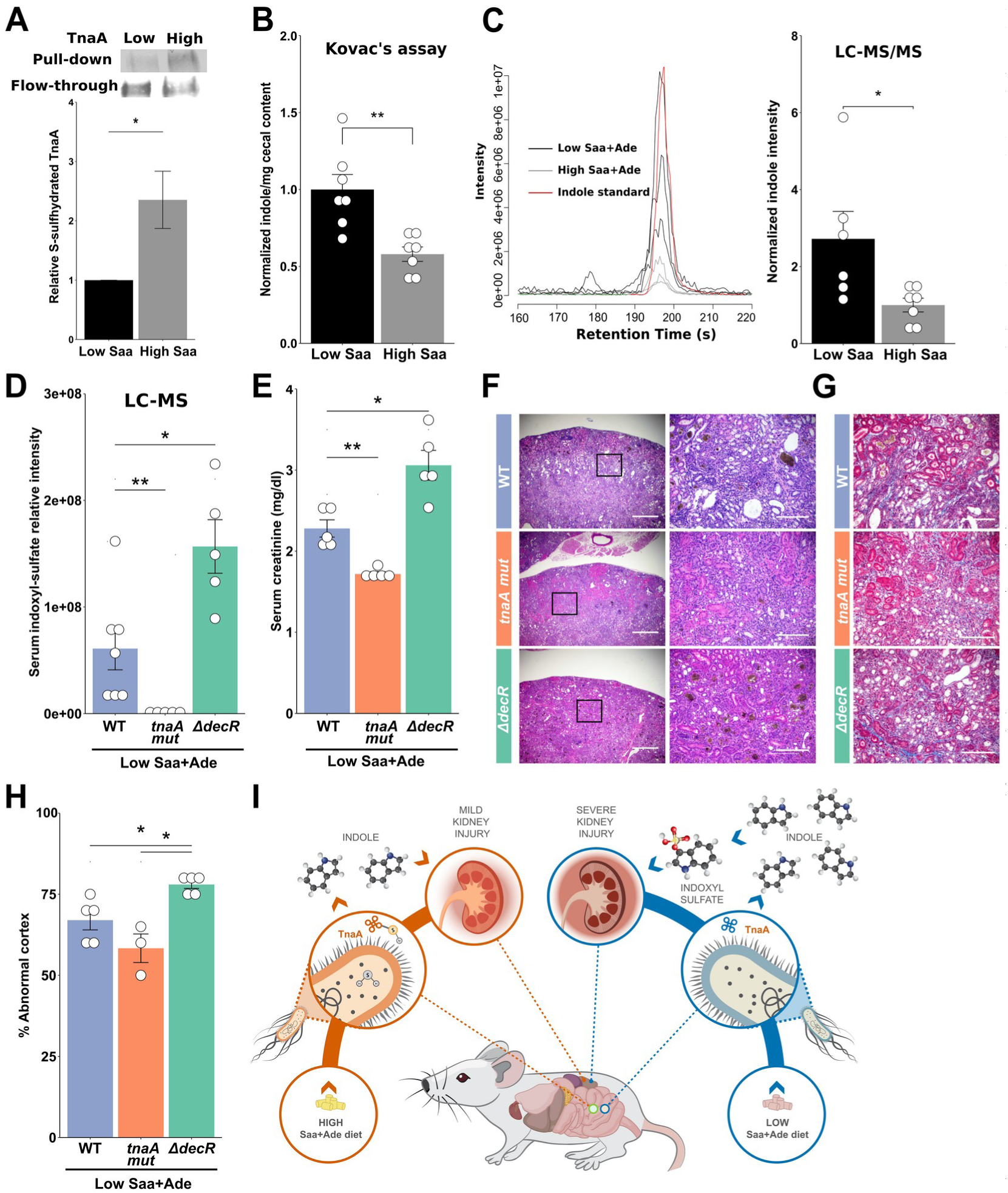
Dietary Saa Modulate Cecal Indole Levels, Serum Indoxyl-Sulfate Levels, and Kidney Function in a Mouse CKD Model. A. Western blot analysis of TnaA of S-sulfhydration pull-down and flow-through samples from cecal contents and **B.** Kovac’s assay measurement of indole levels in cecal contents from ASF*^E. coli^* mice on Saa diets. **C.** LC-MS/MS analysis of indole levels in cecal contents from ASF*^E. coli^* mice on Saa diets. Left, spectra representative of an experiment with 3 mice per group and indole standard. **D.** LC-MS measurements of serum indoxyl-sulfate in ASF mice on low Saa+Ade diet, colonized with *E. coli* strains. **E.** Serum Cre levels in ASF mice colonized with *E. coli* strains on low Saa+Ade diets. **F.** Representative H&E staining and **G.** Representative trichrome staining of kidneys from mice in E. **H.** Histology-based renal injury score. **I.** Illustration showing the effects of low and high Saa-Ade diets on gut microbial activity and the consequences for renal function. Data represent 3 independent experiments for **A**, **B**, **C**, **D**, **E** and **H**. Symbols represent individual mice. Bars represent mean ± SEM. * P value < 0.05, ** P value < 0.01. Mann-Whitney test for **A**, **B** and **C,** two-way ANOVA with Tukey’s post-hoc test for **D**, **E** and **H**.

Herein, we report that sulfide derived from bacterial metabolism of dietary Saa regulates *E. coli* indole production through inhibition of tryptophanase by S-sulfhydration. Our work shows that a dietary component can be metabolized by the microbiota to generate a post-translational modification of microbial proteins that affects host physiology and offers a framework for how host-diet-microbiota interactions can contribute to or stave off to disease states such as CKD (Fig. 4I). While metatranscriptomics have provided a window into functional changes within a community, our results demonstrate that at times we need to delve even more deeply—beyond transcriptomics and even proteomics—to the effect of a single modification on one specific protein. Indeed, our work leverages a subtle dietary change, which does not result in microbial composition changes in our mouse CKD model, to show that production of indoles by *E. coli* is differentially affected by levels of sulfide endogenously produced by gut bacteria. Hence, these results emphasize not only the effect of bacterial metabolism on host physiology, but also potential microbe-microbe interactions driven by bacterial post-translational modifications mediated by host diet beyond the S-sulfhydration studied herein. Diet is a crucial aspect in managing CKD (*22, 23*) and we hypothesize that administration of TnaA inhibitors, such as sulfide donors, may help reduce gut indole levels and thus mitigate kidney damage. In support of this concept and its broad application, other gut bacteria, especially members of the *Bacteriodetes* phylum also encode for TnaA homologs (*24*), and a high degree of homology exists between bacterial TnaA alleles (Fig. S8). Overall, our study elucidates an interaction between diet, microbial metabolism, and kidney function, mediated by post-translational protein regulation. These findings might shed light on managing CKD and provide clinical approaches that target the microbiota and the enzymatic activities of its proteome to improve human health.

## Acknowledgments

The authors would like to thank Jennifer X. Wang (Small Molecule Core Mass-Spectrometry, Harvard University), Bogdan Budnik and Renee Robinson (Harvard Center for Mass Spectrometry Proteomics), Ryan Kuntz and Sanjukta Thakurta (Thermo Fisher Scientific Center for Multiplexed Proteomics, Harvard University), Jessica K. Lang and Kate Rosinski (Harvard T.H. Chan Gnotobiotic Center for Mechanistic Microbiome Studies) for technical support and members of the Garrett lab for their discussion of this work. **Funding:** This work was supported by CRI Irvington post-doctoral fellowship to LL, NIH T32 AI118692 and F31 DK121375 to Y.G.C., and R01CA202704 and R24DK110499 to W.S.G. **Author contributions:** Conceptualization: LL and WSG. Formal analysis: LL. Funding acquisition: WSG. Investigation: LL, YGC, and JNG. Software: LL. Supervision: WSG. Visualization: LL, YGC and WSG. Writing – original draft: LL and WSG. Writing – review and editing: LL, YGC, JNG and WSG. **Competing Interests:** The authors declare no competing financial interests. WSG is on the SAB of Kintai Therapeutics, Leap Therapeutics, Evelo Biosciences, and Tenza Inc. **Data and materials availability:** All data are available in the manuscript or the supplementary materials. Raw 16S rDNA sequences were deposited in NCBI SRA databank under the bioproject accession PRJNA603373.

## Supplemental Material and Methods Mice and dietary interventions

C57BL/6J (B6) mice were obtained from Jackson Laboratory and were housed at the Harvard T.H. Chan School of Public Health (HSPH) barrier facility with constant ambient temperature of 24°C and 12h of day/night cycles. For gnotobiotic experiments, mice were housed at our gnotobiotic facility in semi-rigid isolators and experiments were conducted in individual ventilated cages. Routine qPCR analyses (using universal 16S rDNA primers (*25*)) were performed on fecal samples and cage swabs to validate the gnotobiotic status of the animals. In order to generate mice that harbor the Altered Schaedler Flora (ASF) microbiota, germ-free (GF) mice were gavaged with cecal contents of ASF mice. Colonization was determined by qPCR as previously reported (*26*). Sulfur amino acids (Saa) diets were formulated based on previous literature (*27, 28*) to represent edge cases of Saa consumption, *i.e.* low vs high Saa diets (see Table S1 for the diets formulations) and manufactured by Research Diets, Inc. The lower Saa diet contains enough methionine to avoid the effects of methionine restriction (*9*). 6-8 weeks old SPF or ASF mice were placed on an Saa diet and maintained on it until the experimental end-point. For generating ASF mice colonized with *E. coli* strains, 6-8 weeks old B6 ASF mice were gavaged with 5×10^7^-5×10^8^ colony forming units (CFU) of *E. coli* strains, and cecal colonization was confirmed at the end-point. For the preclinical CKD model, 0.2% adenine was added to the Saa diets and 6-8 weeks old B6 SPF, GF, ASF or ASF*^E. coli^* mice were maintained on Saa+0.2% adenine (Saa+Ade) diets for 7 weeks (*8*). The diets were manufactured by Research Diets, Inc. Animal studies and experiments were approved and carried out in accordance with Harvard Medical School’s Standing Committee on Animals and the National Institutes of Health guidelines for animal use and care.

### Bacterial strains and media

*E. coli* K-12 BW25113 strain were used in all the experiments (H_2_S production, indole production and *in vivo* experiments). For technical reasons, *E. coli* K-12 MG1655 was used in the cloning process to generate *ΔdecR* and *tnaA-his, and E. coli* K-12 W3110 was used for expressing and purifying TnaA-His. Bacteria were grown in LB broth (Merck) at 37°C with shaking (250 rpm) aerobically or without shaking anaerobically and where mentioned, LB was supplemented with L-cysteine (Sigma-Aldrich), sodium hydrosulfide (Sigma-Aldrich) and/or L-tryptophan (Sigma-Aldrich). For selection on LB + Chloramphenicol (Cm) agar plates, a concentration of 10µg/ml Cm was used. The *E. coli tnaA739::kan* strain was obtained from Yale University Coli Genetic Stock Center (CGSC) as part of the Keio collection (*29*).

### Genetic manipulations and cloning of *E. coli*

To generate the in-frame *decR* deletion mutant, 1000 bp up-stream and down-stream of *decR* coding sequence were amplified using 2 consecutive PCR reactions (list of primers, purchased from Sigma-Aldrich, is provided in Table S4) to construct a PCR product that contains only the first and last codons of *decR*. The PCR product was then digested with BamHI and NotI restriction enzymes, ligated into the pKOV plasmid (obtained from AddGene) and chemically transformed into *E. coli* MG1655. Bacteria were plated on LB+Chloramphenicol (Cm) plates and incubated at 30°C, due to the temperature sensitivity of the origin of replication (*30*). The pKOV-*decR* plasmid was extracted from colony PCR positive colonies and was sequenced by Sanger sequencing (Genewiz) to verify the sequence of the insert. *E. coli* BW25113 strain was then transformed with the pKOV-*decR* plasmid and plated on LB+Cm plates at 30°C overnight. Resistant colonies were inoculated into LB+Cm and grown at 42°C overnight to force the integration of the plasmid into the bacterial genome. Cm^R^ colonies were picked and grown in LB without antibiotics for three passages at 30°C to allow the excision of the plasmid. Finally, the bacteria were plated on salt-free LB plates containing sucrose, which should restrict the growth of bacteria that still contain the plasmid (and the *sacB* gene). Colony PCR was used to validated the deletion site. A similar procedure was used to replace the chromosomal *tnaA* copy with a *tnaA-his* allele, except that the initial PCRs were leveraged to sew the his-tag into the *tnaA* PCR product (primer sequences provided in Table S4).

### Protein extraction

Bacterial cultures were grown in LB medium until mid-log growth (OD_600nm_=0.5-0.7) and then bacteria were pelleted by centrifugation at 4000rpm for 10min at 4°C. When indicated, the bacterial cultures where supplemented with 5mM cysteine or 1mM NaHS after 3h of growth, allowed to grow for an additional 1h and then pelleted. The supernatants were removed and the pellets were washed once in cold PBS to remove extracellular proteins. Washed bacterial pellets were resuspended in 1ml of cold lysis buffer (100mM Tris-HCl pH 7.1, 150mM NaCl, 1mM EDTA, 0.5% deoxycholate and 0.5% Triton X-100) and transferred into a 2ml tube containing 300µl zirconium beads (20 micron). The lysis buffer was de-gassed using argon to reduce loss of S-sulfhydration signal by oxidation and supplemented with Complete Ultra proteases inhibitor cocktail (Roche) and Phospho-Stop phosphatase inhibitor (Roche). The tubes were then placed in a bead-beater for 2min and then centrifuged for 10min at 14,000rpm at 4°C. The supernatant was transferred to a new 1.5ml tube and immediately frozen in liquid N_2_ to reduce the possibility of cysteine residues oxidation. Protein quantification was performed using a BCA assay (Thermo Fisher) on a 1:10 diluted sample. For mouse cecal samples, ceca of 3 mice on low or high Saa diet were pooled and resuspended in cold TBS buffer to generate one sample under each condition. The samples were gently vortexed for 10min to homogenize the solution. Then, the samples were centrifuged for 3min at 200g to remove undigested food particles and the supernatant was transferred to a new 1.5ml tube and centrifuged for 10min at 14000rpm at 4°C. The supernatant was discarded and the pellets were resuspended in 0.6ml cold lysis buffer and transferred to a 2ml tube with 300µl of zirconium beads (20micron) and processed similarly to the bacterial samples. For analyzing TnaA-His S-sulfhydration, bacterial cultures of *E. coli tnaA-his* W3110 strain were grown in LB for 3h and then 5mM cysteine was added for 1h. Protein was extracted as mentioned above, purified using the His-Spin protein miniprep (Zymo research) and desalted using Zeba micro-columns (Thermo Fisher) into 100mM TEAB solution. TnaA purity was determined by Coomassie blue staining.

### Pull-down of S-sulfhydrated proteins

Frozen lysates were thawed on ice and 1-2mg of protein in 125µl were incubated with occasional stirring at room temperature for 1h with freshly prepared 100µM maleimide-PEG2-biotin to label the thiol groups. As needed, samples were concentrated using Amicon Ultracel 3K nanosep columns (Millipore). To remove the unbound maleimide molecules, the samples were desalted using Zeba micro-columns (Thermo Fisher) into binding buffer (50mM Tris-HCl pH 7.5, 0.1% SDS, 150mM NaCl, 1mM EDTA and 0.5% Triton X-100). Sample volume was adjusted to 250µl and samples were incubated overnight (16h) at 4°C with 50µl of pre-washed high-capacity binding streptavidin-agarose beads (Thermo Fisher). The following day the samples were moved onto a micro-column (Thermo Fischer) and centrifuged for 1min at 1000g, the flow-through was collected and labeled as flow-through. Then the beads were washed (all washes were 250µl at 1000g for 1 min) on-column 3 times with wash buffer A (50mM Tris-HCl pH 7.5, 150mM NaCl, 0.5% Triton X-100), followed by 3 washes with wash buffer B (50mM Tris-HCl pH 7.5, 600mM NaCl, 0.5% Triton X-100) and finally one wash with elution buffer (50mM Tris-HCl, 150mM NaCl), before incubation with 500µl elution buffer supplemented with 20mM dithiothreitol (DTT) for 30min and then eluted by 1min centrifugation at 1000g. The pull-down samples were concentrated using Amicon Ultracel 3K nanosep (Millipore) by centrifuging at max speed for 10min or until the samples contained 25-30µl. The beads were resuspended in 300µl elution buffer, boiled for 10min and collected as beads-bound fractions. Pull down fractions were visualized by either Coomassie blue (Bio-Rad) or silver stain (Thermo Fisher).

### FASP on-column and TMT labeling

On-column protein digestion and labeling were performed using FASP digestion kit (Expedeon) following the iFASP protocol (*31*). To increase the recovery of peptides, micro-columns (10kDa MWCO) and collection tubes were incubated overnight in 5% Tween-20 and soaked twice (10min each) with sterile double distilled water (DDW). Then the micro-columns were washed twice with 500µl of mass-spec grade water (Roche). 3µl of 200mM TCEP were added to 30µl of pull-down protein sample in 1.5ml tube and incubated for 1h at 55°C. Then, the samples were cooled to room temperature and 200µl of 8M urea (in 0.1M Tris-HCl, pH 8.5) were added. The samples were transferred to 10kDa MWCO micro-columns and spun at 14000g for 15min. The membranes were washed with 200µl urea 8M and then 100µl of 0.05M iodoacetamide in 8M urea solution were added. The tubes were shaken for 1min at 600rpm and then incubated at room temperature in the dark for 20min, before being spun again at 14000rpm for 15min, washed twice with 200µl 8 M urea solution pH 8.5, and washed 3 times with 100µl of 100mM TEAB solution. 75µl of 1:50 dilution of Trypsin (Thermo Fisher) at 1µg/µl in 50mM acetic acid with 0.02% ProteaseMAX (Promega) were added to the micro-columns, which were then incubated at 37°C at 600 rpm for 16h. The tubes were sealed with parafilm to avoid drying of the membranes. Next, TMT reagents (Thermo Fisher) were equilibrated to room temp and dissolved in 41µl of anhydrous acetonitrile for 5min with occasional vortexing. Then, each TMT label was added to a micro-column and incubated at room temperature in the dark at 600rpm for 1h. The reactions were quenched by adding 8µl of 5% hydroxylamine and room temperature incubation for 30min at 600rpm. Labeled peptides were eluted by passing 40µl of 100mM TEAB over the columns 3 times followed by 50µl of 0.5M NaCl solution. The TMT labeled channels (3 repeats of WT *E. coli*, WT *E. coli* without DTT elution and *ΔdecR* lysate, as well as a reference channel, made by combining equal volumes of the 9 samples prior to TMT labeling), were combined into one tube and dried using a speed-vac. The dried pooled sample was resuspended in 1ml 1% triflouroacetic acid (TFA) solution and incubated for 30min with shaking at 600rpm and then dried again using a speed-vac. The pooled sample was then resuspended in 300µl of 0.1% TFA and fractionated using the Pierce High pH Reversed-Phase Peptide Fractionation Kit (Thermo Fisher) into 5 fractions (10%, 15%, 20%, 25% and 50% acetonitrile), dried in a speed-vac and resuspended in 0.1% formic acid.

### Metabolite extractions for LC-MS/MS analysis of indole and indoxyl-sulfate

Extraction of metabolites from cecal samples was performed similar to (Jin et al., 2014; Sellick et al., 2010) (*32, 33*) by collecting cecal contents into empty pre-weighed 2ml tubes; and after weighing the contents, 1.5ml of cold methanol/chloroform (2:1 v/v) solution were added. The samples were vortexed and homogenized with a wide-bore tip on ice and then centrifuged for 10min at 15,000g at 4°C. The supernatant was transferred to a new 5ml tube, 0.6ml of ice-cold double-distilled water were added and the samples were vortexed and centrifuged at 15,000g for 5min at 4°C to obtain phase separation. The upper aquatic phase and lower organic phase were collected carefully without dispersing the proteinaceous interface into 1.5ml tubes and kept in −80°C until LC-MS/MS analysis. For bacterial culture indole measurements, overnight bacterial cultures were diluted 1:100 and grown for 3h at 37°C in LB, then either mock, 5mM L-cysteine or 1mM NaHS were added for 1h. Bacterial cultures were harvested by centrifugation, the supernatants were filter-sterilized through 0.2µm filters and 225µl of supernatant were transferred to a new 1.5ml tube. Then, 25µl of 10mM L-tryptophan was added and the samples were incubated 1h at 37°C, before 250µl of 20% TCA were added and the samples were incubated on ice for 15min to precipitate proteins. The samples were then centrifuged for 10min at 4°C and the supernatants were kept at −80°C until LC-MS/MS analysis. For serum samples, mouse blood was collected into serum separator tubes (BD) tubes, inverted 5 times and allowed to clot for 30min at room temperature. Then, samples were centrifuged for 15min at 1300g at 4°C and the serum layer was carefully removed into a new 1.5ml tube without disturbing the buffy coat layer. The samples volume was adjusted to 160µl with PBS, 40µl of trichloroacetic acid (TCA) were added to 20% final TCA concentration and samples were incubated on ice for 15min to precipitate proteins. The samples were then centrifuged at max speed for 10min at 4°C and the supernatants were transferred to new 1.5ml tubes and kept at −80°C until LC-MS/MS analysis.

### Mass spectrometry analysis

The TMT fractions were analyzed by LC-MS^3^ on an Orbitrap Fusion™ Lumos™ Tribrid™ mass spectrometer at the Thermo Fisher Scientific Center for Multiplexed Proteomics (TCMP) at Harvard Medical School. Labeled peptide samples were analyzed with an LC-MS^3^ data collection strategy (McAlister GC et al. (2014) Anal. Chem. 86:7150-8) on an Orbitrap Fusion mass spectrometer (Thermo Fisher Scientific) equipped with a Thermo Easy-nLC 1200 for online sample handling and peptide separations. Resuspended peptide from previous step was loaded onto a 100µm inner diameter fused-silica micro capillary with a needle tip pulled to an internal diameter less than 5µm. The column was packed in-house to a length of 35cm with a C_18_ reverse phase resin (GP118 resin 1.8μm, 120Å, Sepax Technologies). The peptides were separated using a 180min linear gradient from 6% to 35% buffer B (90% ACN + 0.1% formic acid) equilibrated with buffer A (5% ACN + 0.1% formic acid) at a flow rate of 500 nL/min across the column. The scan sequence for the Fusion Orbitrap began with an MS1 spectrum (Orbitrap analysis, resolution 120,000, scan range of 350 - 1350m/z, AGC target 1 x 10^6^, maximum injection time 100ms, dynamic exclusion of 60 seconds). The “Top10” precursors was selected for MS2 analysis, which consisted of CID (quadrupole isolation set at 0.5Da and ion trap analysis, AGC 2.5×10^4^, Collision Energy 35%, maximum injection time 150ms). The top ten precursors from each MS2 scan were selected for MS3 analysis (synchronous precursor selection), in which precursors were fragmented by HCD prior to Orbitrap analysis (Collision Energy 55%, max. AGC 2×10^5^, maximum injection time 150ms, resolution 50,000, and isolation window set to 1.2 – 0.8). *E. coli* TnaA-His samples were analyzed at the Mass Spectrometry and Proteomics Resource Laboratory at Harvard University. TnaA-His was not reduced and/or alkylated to preserve the state of native cysteine PTMs. LC-MS/MS was performed on a Orbitrap Elite™ Hybrid Ion Trap-Orbitrap Mass Spectrometer (Thermo Fischer, San Jose, CA) equipped with Waters Aquity nano-HPLC. Peptides were separated onto a 100µm inner diameter microcapillary trapping column packed first with approximately 5cm of C18 Reprosil resin (5µm, 100Å, Dr. Maisch GmbH, Germany) followed by analytical column ∼20cm of Reprosil resin (1.8µm, 200Å, Dr. Maisch GmbH, Germany). Separation was achieved through applying a gradient from 5-27% ACN in 0.1% formic acid over 90min at 200nl/ min. Electrospray ionization was enabled through applying a voltage of 1.8kV using a home-made electrode junction at the end of the microcapillary column and sprayed from fused silica pico tips (New Objective, MA). The mass spectrometry survey scan was performed in the Orbitrap in the range of 395–1,800m/z at a resolution of 6×10^4^, followed by the selection of the twenty most intense ions (TOP20) for CID-MS2 fragmentation in the Ion trap using a precursor isolation width window of 2m/z, AGC setting of 10,000, and a maximum ion accumulation of 200ms. Singly charged ion species were not subjected to CID fragmentation. Normalized collision energy was set to 35V and an activation time of 10ms. Ions in a 10ppm m/z window around ions selected for MS2 were excluded from further selection for fragmentation for 60s. The same TOP20 ions were subjected to HCD MS2 event in Orbitrap part of the instrument. The fragment ion isolation width was set to 0.7 m/z, AGC was set to 50,000, the maximum ion time was 200ms, normalized collision energy was set to 27V and an activation time of 1ms for each HCD MS2 scan. Metabolite samples were analyzed for indole and indoxyl sulfate content at the Small Molecule Mass Spectrometry core at Harvard University. Quantification of indole by LC/MS-MS were carried out on a Thermo Scientific Dionex UltiMate 3000 UHPLC coupled to a Thermo Q Exactive Plus mass spectrometer system (Thermo Fisher Scientific Inc, Waltham, MA) equipped with an APCI probe for the Ion Max API source. Data were acquired with Chromeleon Xpress software for UHPLC and Thermo Xcalibur software version 3.0.63 for mass spectrometry and processed with Thermo Xcalibur Qual Browser software version 4.0.27.19. 3µL sample was injected onto the UHPLC including an HPG-3400RS binary pump with a built-in vacuum degasser and a thermostated WPS-3000TRS high performance autosampler. An Xterra MS C18 analytical column (2.1×50mm, 3.5µm) from Waters Corporation (Milford, MA) was used at the flow rate of 0.3mL/min using 0.1% formic acid in water as mobile phase A and 0.1% formic acid in methanol as mobile phase B. The column temperature was maintained at room temperature. The following gradient was applied: 0-6min: 20-100% B, 6-8min: 100%B isocratic, 8-8.1min: 100-30% B, 8.1-11.1min, 20%B isocratic. The MS conditions were as follows: positive ionization mode; PRM with the precursor → product ion PRM transition, m/z 118.0651 ([M+H]^+^) → 91.0542 ([C7H7]^+^); normalized collision energy (NCE), 105; resolution, 70,000; AGC target, 2e5; maximum IT, 220ms; isolation window, 1.8m/z; spray voltage, 5000V; capillary temperature, 250°C; sheath gas, 28; Aux gas, 5; probe heater temperature, 363°C; S-Lens RF level, 55.00. A mass window of ±5 ppm was used to extract the ion. Indole was considered detected when the mass accuracy was less than 5ppm and there was a match of isotopic pattern between the observed and the theoretical ones and a match of retention time between those in real samples and the standard. Isotope labeled (^13^C) and native standards of indole and indoxyl sulfate were obtained from Toronto Research Chemicals.

### Computational analysis of mass-spectrometry data

Thermo Fisher RAW files were converted to mzML files using ProteoWizard (*34*). The MS/MS spectra were searched against a target and decoy database comprising the Uniprot *E. coli* K-12 proteome and a list of frequent mass-spectrometry contaminants using MSGF+ (v2017.08.23) (*35*) with the following parameters-protocol 4-t 10ppm-mod MSGF_mod.txt-tda 0-addFeatures 1-maxCharge 4. The modification file included static alkylation of cysteine (57.02146 Da), static TMT labeling of lysine residues and N-termini of peptides (229.162932 Da), and variable oxidation of methionine (15.99491 Da). Post-search peptide filtering was performed using Percolator (v3.1.2) (*36*) and the output psms files was manually filtered to include only psms with q value <= 0.01 and pep score <= 0.05. The filtered psms file was converted to pepXML using OpenMS (v2.2.0) IdfileConverter (*37*) and then TMT MS^3^ reporter ion quantification was performed using pyQuant (v2.1) (*38*). Finally, peptides that had an intensity value of 0 in the reference channel or had a non-zero value only in the reference channel were removed, and proteins that were identified by only one peptide or mapped to contaminants were discarded. Statistical analysis was performed using Kruskal-Wallis and 5% false discovery rate (FDR) in R. For the analysis of TnaA S-sulfhydration, raw data were submitted for analysis in Proteome Discoverer 2.2 (Thermo Scientific) software. Assignment of MS/MS spectra was performed using the Sequest HT algorithm by searching the data against a protein sequence database including all entries from the *E. coli* proteome database as well as other known contaminants such as human keratins and common lab contaminants. Sequest HT searches were performed using a 20ppm precursor ion tolerance and requiring each peptides N-/C termini to adhere with Trypsin protease specificity, while allowing up to two missed cleavages. A MS2 spectra assignment false discovery rate (FDR) of 1% on both protein and peptide level was achieved by applying the target-decoy database search. Visualization of peptide-match spectra was performed using SearchGUI (v3.3.15) (*39*) and PeptideShaker (v1.16.40) (*40*). Indole and indoxyl sulfate analyses were performed using the xcms (3.8.1) package in R.

### Western blot analyses

Equal volumes of pull-down or flow-through samples were incubated at 70°C for 15min with loading buffer. Samples were run on 10% Mini-PROTEAN® TGX™ Precast Protein Gels (Bio-Rad) with Chameleon® Duo Pre-stained Protein Ladder (LiCOr) in Tris-Glycine-SDS run buffer. Proteins were then transferred to an Amersham Protran 0.45µm nitrocellulose membrane in a Tris-Glycine transfer buffer for 1h at 20V at room temperature. The membranes were blocked using 1:2 dilution of Odyssey Blocking PBS Buffer (LiCOr) in PBS for 1h at room temperature. Membranes were then incubated with primary antibodies in blocking buffer + 0.2% Tween-20 overnight. Following 5 washes of PBS+0.2% Tween-20 (5 min each), the membranes were incubated for 1h in blocking buffer+0.2% Tween-20 with secondary antibodies conjugated to a fluorophore (LiCOr). The membranes were washed again for 5 times with PBS+0.2% Tween-20 and then 2 more times with PBS, before being imaged on a LiCOr Odyssey CLx machine. Images were analyzed and quantified using ImageJ2 software (*41*).

### Colorimetric indole measurement using Kovac’s reagent

*E. coli* cultures were grown in 10ml LB at 37°C at 250rpm for 3h and then, for the treatment groups, 5mM L-cysteine or 1mM NaHS were added and the cultures were allowed to grow for an additional 1h before cells were harvested by centrifugation at 14000rpm for 10min at room temperature. Then the supernatants were filter sterilized through 0.2µm filters and 1mM of L-tryptophan was added, and the supernatants were incubated at 37°C for 1h. Then 250µl of 20% w/v TCA was added to 250µl of supernatant and kept on ice for 15min to precipitate proteins. The samples were centrifuged at 14000rpm for 10min, the supernatant was moved to a new 1.5ml tube and 500µl of Kovac’s reagent (Sigma-Aldrich) were added. The samples were vortexed and incubated at 37°C for 30 min, before the top 200µl layer was moved to a 96-well plate, and OD_530nm_ was read. Indole (Sigma-Aldrich) serial dilutions were analyzed for generation of a standard curve. For cecal samples, cecal content was collected into pre-weighed 1.5ml tubes containing 750µl 70% EtOH. The samples were homogenized using vortexing and wide-bore tips and then incubated at 70°C for 10min. After 20min centrifugation at max speed at 4°C, 150µl of supernatant were added to 150µl of Kovac’s reagent, incubated for 30min at room temperature and absorbance at OD_530nm_ was measured.

### *In vitro* Tryptophanase Assays

*E. coli* apo-tryptophanase (Sigma-Aldrich) was resuspended in 1ml 100mM potassium phosphate pH 8 buffer, aliquoted and kept at −20°C. 5µl of apo-tryptophanase were added to 12µl of 100mM potassium phosphate pH 8 buffer with 1mM pyrodxal-5-phosphate (PLP) and various treatments (NaCl, NaHS, L-cysteine, DTT or Na_2_S_4_) and incubated for 45min at 37°C. Then 12 µl of 100 mM potassium phosphate with 5mM L-tryptophan were added and the samples were incubated for 1h at 37°C. Then 250µl of 20% TCA were added to the samples, followed by a 15min incubation on ice to precipitate TnaA. Samples were centrifuged for 10min at max speed, the supernatant was transferred to a new 1.5ml tube and 500µl of Kovac’s reagent were added, followed by vortex and 30min room temperature incubation, before absorbance of the top layer was measured at OD_530nm_.

### Serum creatinine measurements

Mouse serum was extracted as mentioned above and creatinine levels were measured using the Serum Creatinine Colorimetric Assay Kit (Cayman Chemical) in duplicates with a standard curve, following the manufacturer’s protocol.

### Cecal DNA extraction and RT-qPCR analysis

Mouse cecal contents were collected into 1.5ml tube and flash frozen. Upon thawing, cecal contents were resuspended in lysis buffer (100mM Tris-HCl pH 8.0, 15mM EDTA and 2% SDS) and transferred to a 2ml tube containing 300µl zirconium beads (20 micron) and 500µl of TE-saturated phenol (Sigma-Aldrich). The tubes were then placed in a bead-beater for 2min and centrifuged for 10min at 14,000rpm at 4°C. The aqueous phase was transferred to a new 1.5ml tube and an equal volume of phenol:chloroform solution was added. The samples were vortexed and centrifuged for 2 min at max speed at room temperature. This process was repeated 2 more times. Then the aqueous phase was moved to a new tube and 2 volumes of 100% EtOH and 1/10 volume of NaOAc pH 5.2 were added. The tubes were inverted several times and incubated at −20°C for 1 h, before being centrifuged for 20 min at max speed at 4°C, and washed once in cold 70% EtOH. Finally, after air-drying, the samples were resuspended with 100µl of sterile water and DNA concentration was determined using a photospectrometer machine. For RT-PCR analysis, 50 ng of cecal DNA were taken to a reaction with 10µl of SYBR green (KAPA SYBER FAST) and the appropriate primers (Table S4). RT-PCR analysis was performed on a Applied Biosystems Stratagene MX3005P machine. The relative abundance of each ASF bacterium was determined by 2^-ΔCt = (ASF bacterium Ct – Total 16S rRNA Ct)^.

### Bacterial 16S rDNA amplicon library generation and sequencing

The procedures in this section were done in a biological hood to minimize potential contamination and were based on the Earth Microbiome Project protocol (*42*). 50ng of cecal contents extracted DNA were taken to a PCR reaction using the Thermo Fisher Platinum Hot Start PCR Master Mix (cat. no. 13000014) according to the reagent protocol. Forward (10µM) and reverse (1.3µM) primers were used (see Table S4 for primer sequences) to amplify the V4 region of the 16S rRNA gene. For each sample the reverse primer contains a unique 12 bp Golay barcode. Each sample was amplified in triplicate with a 25µl reaction volume per reaction in 96-well plates. Sterile water and *E. coli* genomic DNA served as negative and positive controls, respectively. The PCR reaction started with 3min of 94°C, then 35 cycles of 45s 94°C, 60s 50°C and 90s 72°C, followed by 10min of 72°C. After amplification, the triplicate reactions were pooled and amplicons were purified using AMPure magnetic beads. DNA concentration was determined by dsDNA broad range assay kit (Thermo Fisher) and a sample of several libraries was run on an agarose gel to visualize the specific amplicon. The libraries were pooled so that the final DNA concentration was 50ng/µl and each library had an equal abundance. DNA sequencing was performed on an Illumina Mi-Seq machine at the bio-polymer core of HMS using the Mi-Seq V2 kit with a 250bp paired-end reads. Raw sequences were deposited in NCBI SRA databank under the bioproject accession PRJNA603373.

### Analysis of Bacterial 16S rRNA gene amplicon sequences

Our 16S rRNA gene amplicon sequence analysis was based on the suggested standard operating protocol by Langille et al (*43*) using the microbiome-helper wrapper. Briefly, fastq files were obtained for each library with a median read count of 73,905 250-bp paired-end reads and quality of reads was checked using FastQC (v0.11.5). According to the quality report, the reads were trimmed using the fastx-toolkit to keep only high-confidence base calls. Reads were stitched using PEAR (*44*), converted to fasta format and chimeric reads were filtered out using VSEARCH (*45*). Operational taxonomic units (OTUs) were picked using QIIME V1.9 (*46*) using the sortmerna program (*47*) and OTUs with fewer than 0.1% of the reads were excluded as low-confidence OTUs. Finally, the number of reads in each library was rarefied to the lowest library size. α-diversity and β-diversity (weighted Unifrac PCOA) analyses were conducted using the phyloseq R package v1.30 (*48*) on Rstudio v1.25, as well as visualization of taxonomic compositions. Specific OTU differences between the two diets were analyzed using the phyloseq R package, LefSe (*49*) and MaAsLin (*50*) algorithms using caging as a confounding variable. The python program STAMP was also used to visualize and analyze data (*51*).

### Meta-analysis of CKD patients stool microbiome datasets

Sequence data of 16S rRNA gene amplicon sequencing of CKD patients stool samples (Xu et al., 2017 and unpublished) was downloaded from NCBI (accessions PRJEB9365 and PRJEB5761, respectively). 16S rRNA gene amplicon data was processed as described above. Metagenomic data of unpublished CKD patient stool samples were downloaded from NCBI (accession PRJNA449784) and analyzed by FastQC (v0.11.5), followed by trimming low quality reads and reads that map to the human genome using KneadData (v0.7.2). Next, the filtered reads were used as input for the HUManN2 program (v2.8.1) (*52*), yielding 3 matrices of gene families, metabolic pathways coverage and metabolic pathways abundance, stratified by bacterial species. For *E. coli* abundance analysis, the mean sum-normalized precentage of reads mapping to an *E. coli* gene was calculated and patients without *E. coli* mapped reads were removed from the analysis and then square root transformation was used to normalize the data. PhyloChip data (*12*) was kindly provided by Dr. Vaziri and analyzed using R and the EnhancedVolcano package (v1.4.0).

### H_2_S measurements

Lead acetate paper was prepared by incubating pre-cut Whatman paper in 20mM lead acetate solution for 20min at room temperature and then was dried for 20min at 110°C and kept in the dark (*10*). For bacterial cultures H_2_S production, overnight cultures of *E. coli* strains were diluted to OD_600nm_ of 0.05 in 200µl of LB supplemented with various concentration of L-cysteine in a 96-well plate in triplicate. The plate was tightly covered with lead acetate paper and incubated at 37°C for 7h. The lead acetate papers were then scanned and densitometry analysis was performed using ImageJ2 (*41*). For cecal contents, mouse cecal content was collected into pre-weighed 1.5ml tubes with 300µl of PBS, weighed, homogenized by vortex and pipetting. 200µl were plated in 96-well plates in duplicate and analyzed by lead acetate sulfide assay as described above. For the colorimetric detection of sulfide by the methylene blue method (*53, 54*), cecal contents were collected directly into pre-weighed 1.5ml tubes with 200µl of 1N NaOH to trap free sulfide ions in the S^2-^ non-volatile form, on ice. The samples were homogenized with a wide bore pipette tip (10-20% cecal slurry). The samples were centrifuged at max speed at 4°C for 10min and in parallel a H_2_S standard curve in 1N NaOH was prepared. After centrifugation, 150µl of supernatant were transferred to a new tube and 200µl of DPD/FeCl_3_ reagent (43mM N,N-dimethyl-p-phenylenediamine sulfate and 148mM FeCl_3_ in 4.2M HCl) were added. The samples were vortexed briefly and incubated for 20min at 37°C, thereafter they were centrifuged for 4min at max speed and the supernatants were transferred to a 96-well plate to determine absorbance at OD_670nm_. The pellet from the initial centrifugation was also used to determine bound sulfide levels, as it was washed with 1N NaOH and then resuspended in 200µl of 2% zinc acetate solution, pH 6. Then 300µl of DPD/FeCl_3_ were added and the samples were incubated at 37°C for 20min, centrifuged for 4min at max speed and absorbance was measured in OD_670nm_. For bacterial H_2_S detection, cultures were grown in LB broth for 3h, then L-cysteine was added at 5mM final concentration. After 1h, cultures were centrifuged and 100µl of supernatant were added to 900µl of buffer (100mM potassium phosphate pH 8 and 2.5mM DTT). Then 200µl of DPD/FeCl_3_ were added and the samples were vortexed and incubated for 20min at 37°C before absorbance was read at OD_670nm_.

### Histology

After sacrifice, kidneys were surgically removed from mice and fixed in 4% paraformaldehyde (PFA), embedded in paraffin, sectioned to 5µm and subsequently stained with H&E or Masson’s trichrome reagents. Histological analysis was performed in a blinded fashion by JNG. Abnormal parenchyma was recognized be the presence of one or more of the following: tubular inflammation (tubulitis), tubular dilatation or dropout, interstitial inflammation and or fibrosis. The extent of crystal deposition was also noted. Quantitative scoring was performed as follows: The extent of abnormal (inflamed) renal parenchyma was visually estimated as a percentage of the total cortical area, in well-oriented sections which included both the renal cortex and medulla.

## Statistical analyses

All the statistical analyses were performed using R (v3.4-3.6) on RStudio (v1.25). List of packages used is provided in Table S5. Mann-Whitney and Kruskal-Wallis were performed as the default statistical tests, unless mentioned otherwise in the figure legends.

---

**Fig S1.**
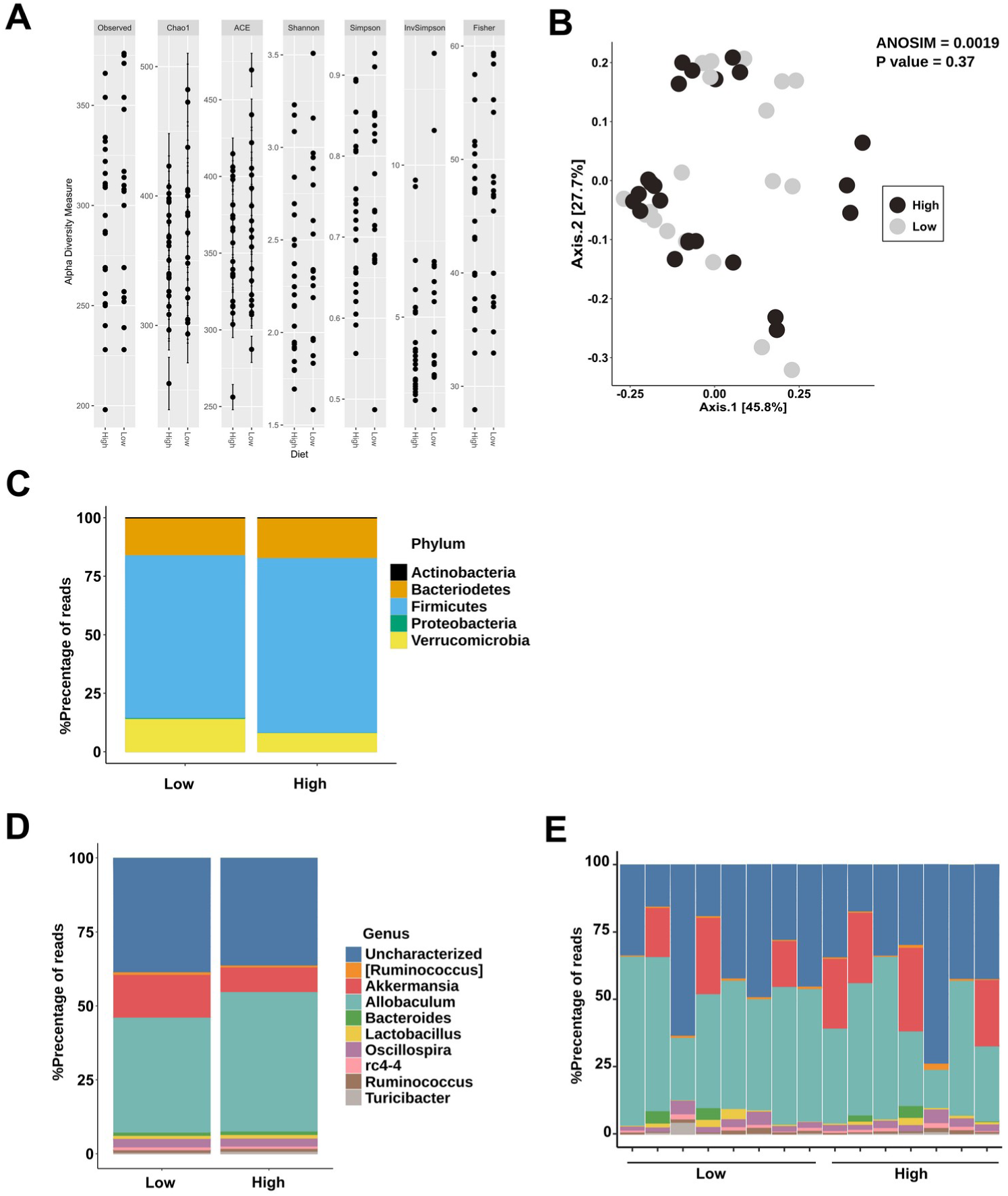
Cecal microbiome 16S rRNA gene amplicon analysis of SPF mice on Saa diets. A. Alpha-diversity measures. **B.** Beta-diversity, weighted UniFrac analysis. **C.** Relative abundances of bacterial phyla. **D.** Relative abundances of the top 10 bacterial genera. **E.** Same analysis as in **D**, each X-axis tick represents a cage, allowing for observation of caging effects. n = 19 mice on low Saa diet and 24 mice on high Saa diet.

**Fig S2.**
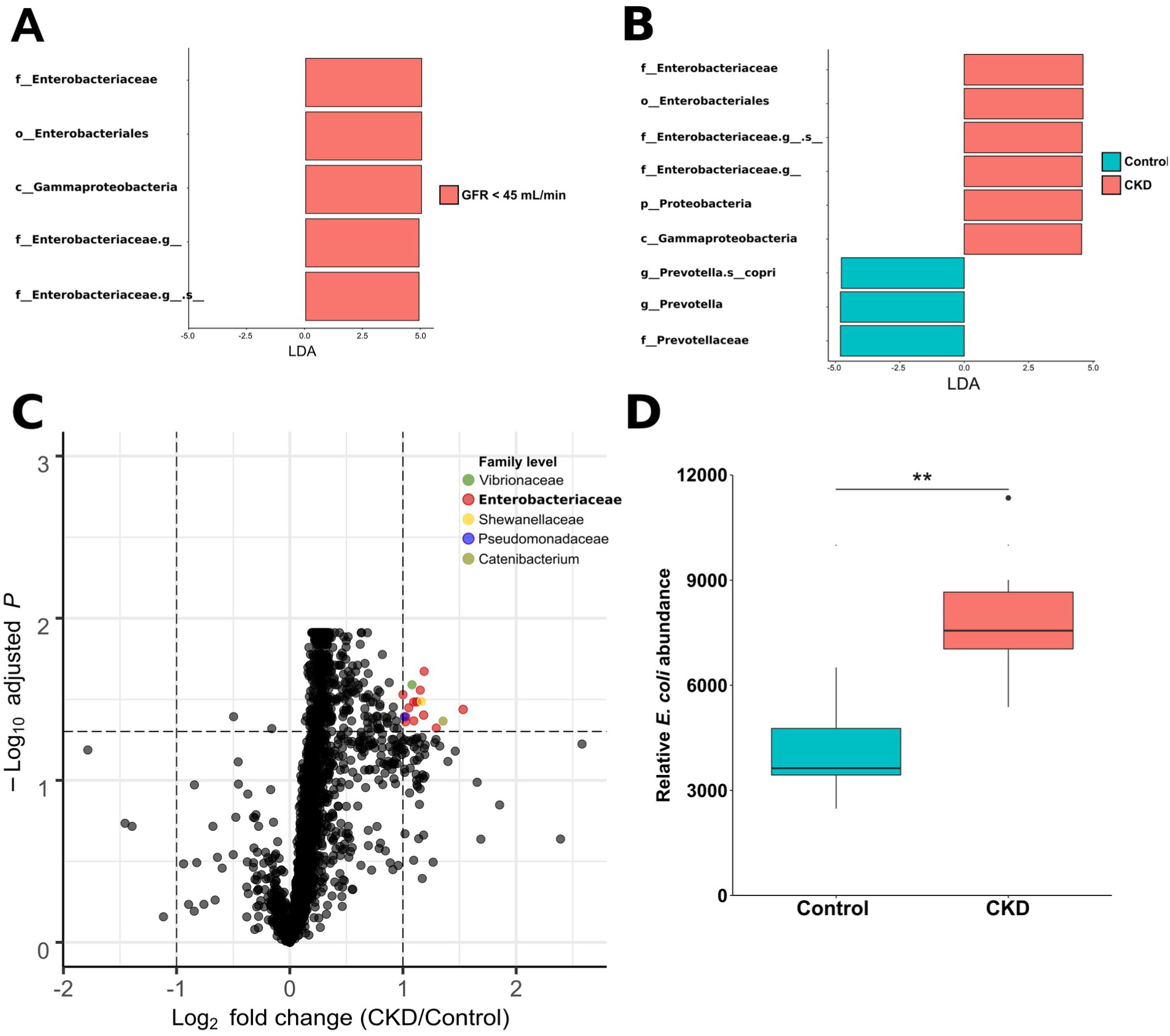
CKD Patient Fecal Microbiome Data Analysis Reveals *Enterobacteriaceae* Enrichment. A. LEfSe analysis of 16S rRNA gene amplicon survey data from Xu *et al.* 2017. **B.** LEfSe analysis of 16S rRNA gene amplicon survey data from an unpublished CKD patient cohort (NCBI accession PRJEB5761). For clarity, taxonomy is shown from the class level. **C.** Volcano plot of PhyloChip analysis data from Vaziri *et al.* (2013). Taxa with fold change > 2 and q-value < 0.05 are labeled in color with their family level taxonomy. **D.** Boxplot representation of the combined averaged relative abundance of 7 *E. coli* strains measured in the fecal samples from CKD and non-CKD subjects using PhyloChip analysis. * P value < 0.05, ** P value < 0.01.

**Fig S3.**
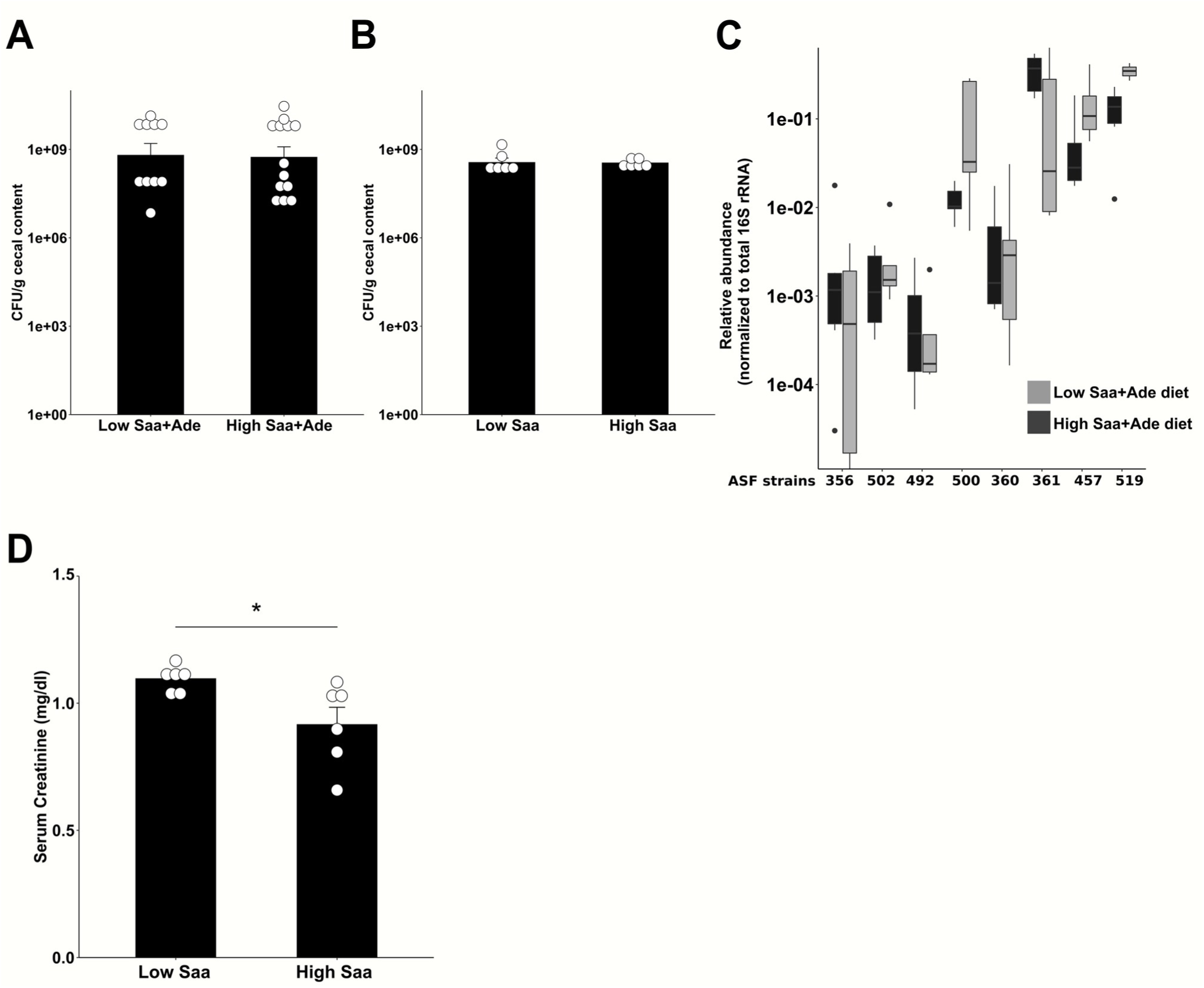
Supplementary data for Fig 1. **A.** Colonization of ASF mice with *E. coli* on Saa+Ade diets. **B.** Colonization of ASF mice with *E. coli* on Saa diets. **C.** Relative abundances of ASF strains in cecal contents of mice on Saa+Ade diets. **D.** Serum Cre levels from WT ASF*^E. coli^* mice on low vs. high Saa diets. Data represent 3 independent experiments for **A**, **B**, **C** and **D**. Symbols represent individual mice. Bars represent mean ± SEM. * P value < 0.05. Mann-Whitney test for **D**.

**Fig S4.**
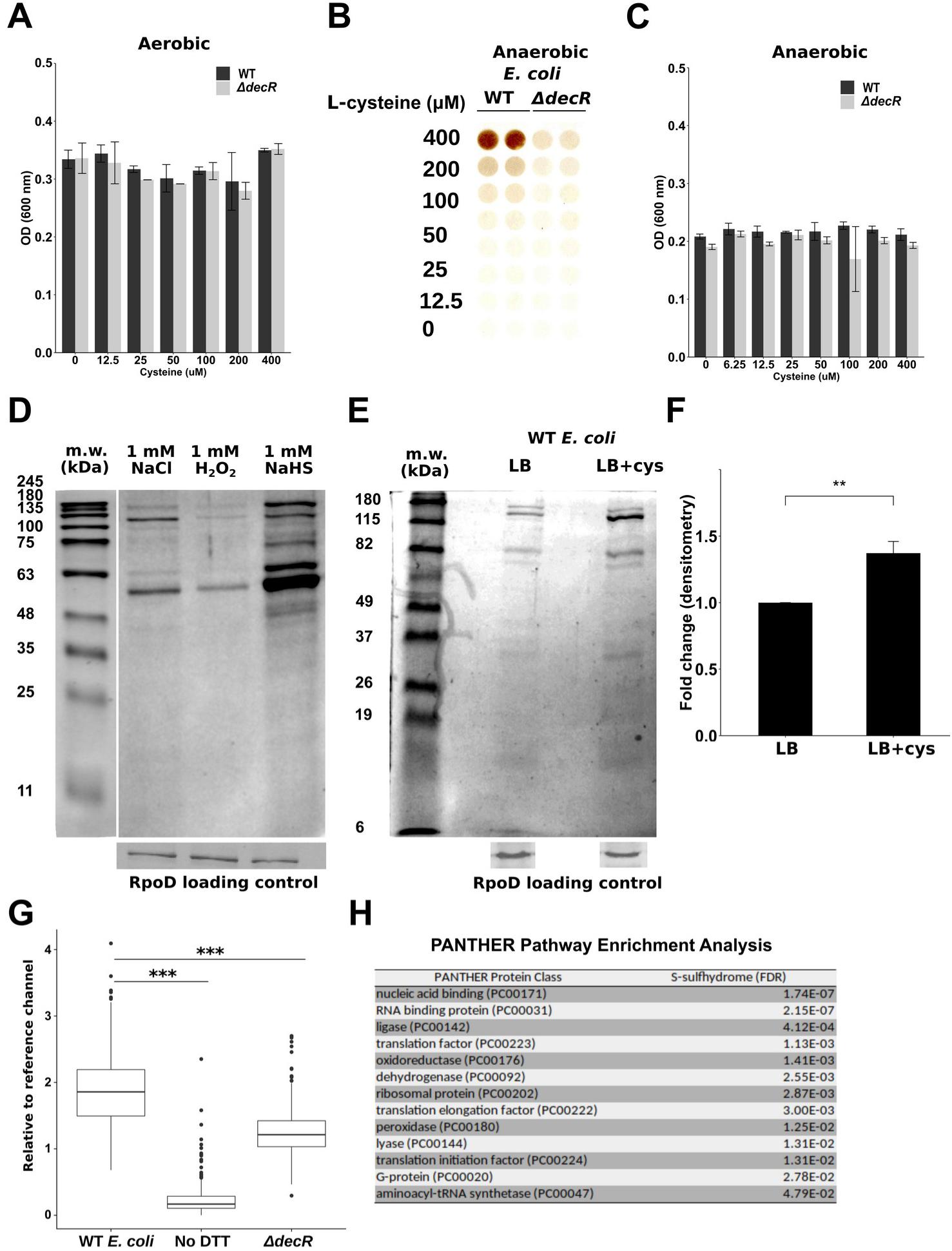
Supplementary data for Fig 2. **A.** Final OD_600_ of WT and Δ*decR E. coli* cultures grown in LB supplemented with cysteine under aerobic conditions. **B.** Lead acetate detection of H2S production by WT and Δ*decR E. coli* cultures grown in LB supplemented with cysteine under anaerobic conditions. **C.** Final OD_600_ of WT and Δ*decR E. coli* cultures grown in LB supplemented with cysteine under anaerobic conditions. **D.** Coomassie stain of S-sulfhydrated proteins from WT *E. coli* lysates treated with NaCl, H2O2 or NaHS. Lower gel shows Western blotting of RpoD in the flow-through samples, as loading control. Data are representative of 3 independent experiments. **E.** Coomassie stain of S-sulfhydrated proteins from WT *E. coli* grown in LB or LB supplemented with 0.4mM cysteine. Lower gel shows Western blotting of RpoD in the flow-through samples, as loading control. **F.** Quantification of Coomassie stains from E. **G.** Boxplot representation of the data presented in Fig. 2F**. H.** Pathway enrichment analysis using the PANTHER database (Mi et al., 2019) of the 212 S-sulfhydrated proteins. Pathways with q-value < 0.05 are reported. Data are representative of 3 independent experiments for **A**, **B**, **C**, **D**, **F** and **G**. Bars represent the mean ± SEM. ** P value < 0.01, *** P value < 0.001. Mann-Whitney test for **F**.

**Fig S5.**
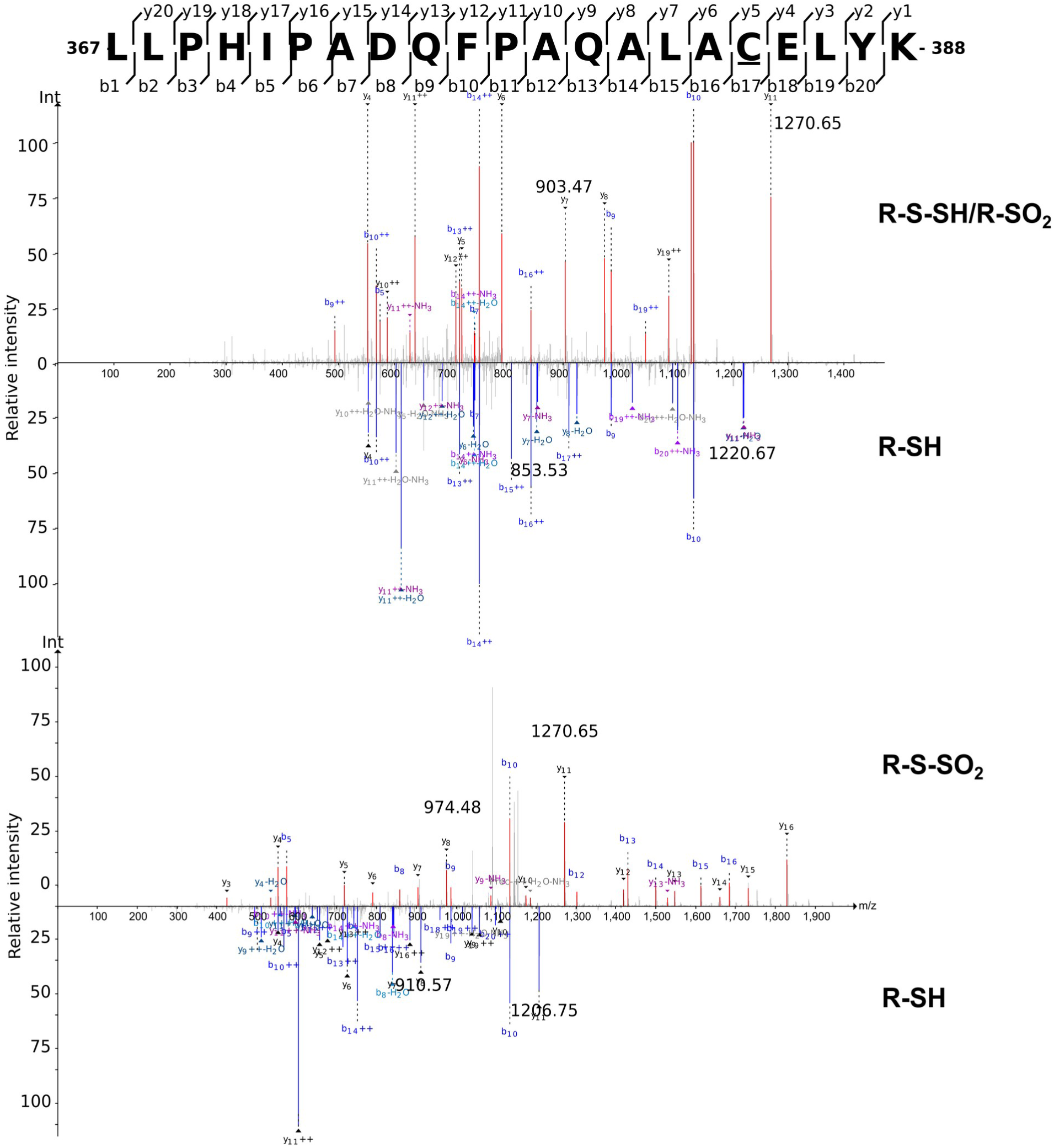
LC-MS/MS m/z spectra of a TnaA S-sulfhydrated peptide. The top spectrum shows the addition of +32 Da on C363, red series ions represent the native cysteine residue (R-SH) peptide and the blue series ions represent the R-SO2/R-S-SH cysteine modification. The lower spectrum represents the addition of +64 Da on C363, red series ions represent the native cystine residue (R-SH) and the blue series ions represent the oxidized S-sulfhydrated (R-S-SO2) or the polysulfhydrated (R-S-S-S) cysteine residue.

**Fig S6.**
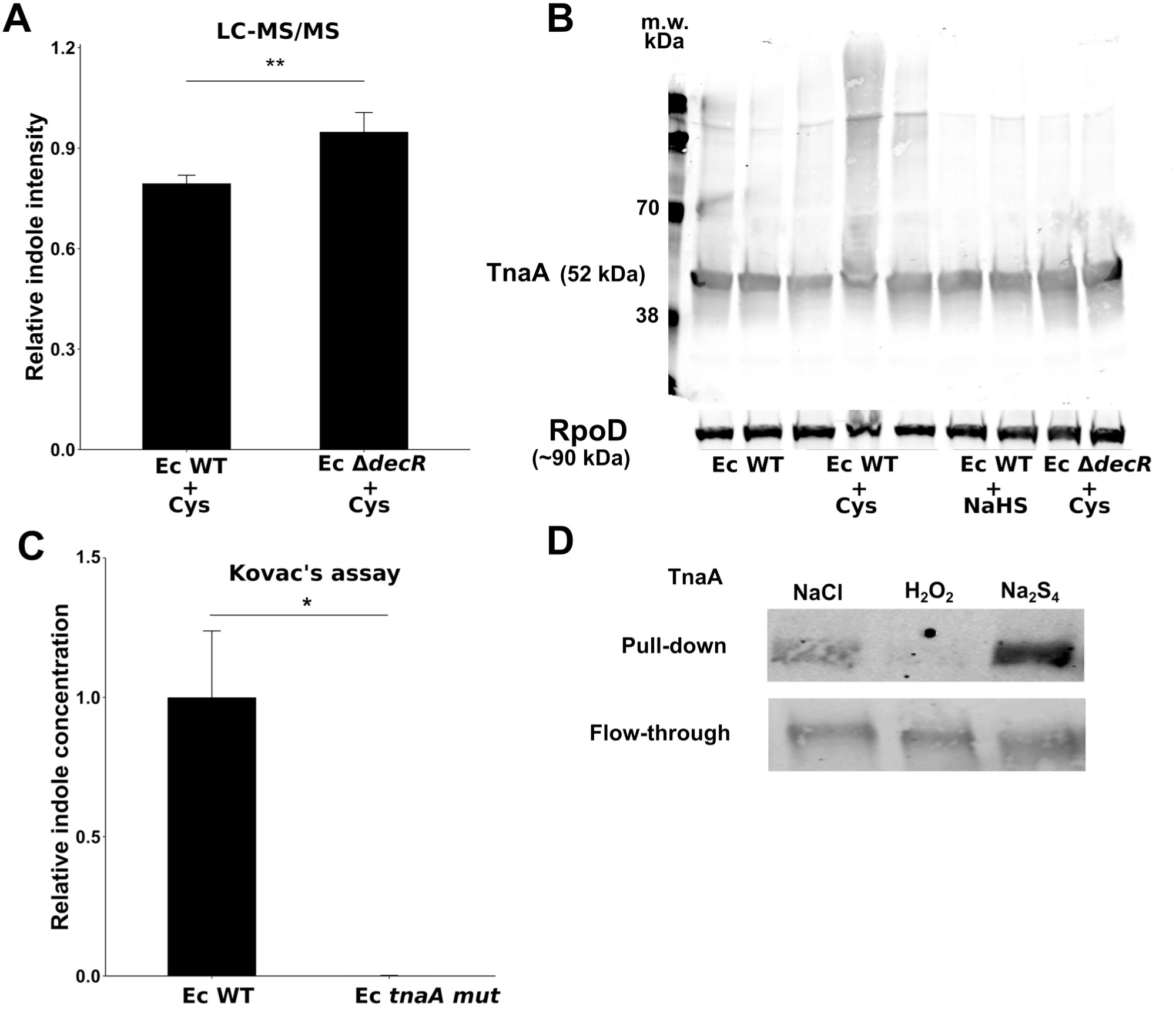
Supplementary data for Fig 3. **A.** Relative indole levels of WT and Δ*decR E. coli* cultures grown aerobically in LB supplemented with cysteine measured by LC-MS/MS. **B.** Western blotting of TnaA in WT *E. coli* cultures grown aerobically in LB or LB supplemented with cysteine or NaHS, or Δ *decR E. coli* grown in LB. Lower gel shows Western blotting for RpoD as loading control. **C.** Relative indole levels of WT and *tnaA mut E. coli* cultures grown aerobically in LB measured by Kovac’s assay. **D.** Western blotting of the S-sulfhydration pull-down fractions of purified *E. coli* TnaA treated with NaCl, H2O2 or Na2S4. Flow through samples represent loading control. Data represent 3 independent experiments for **A**, **B**, **C** and **D**. Bars represent mean ± SEM. * P value < 0.05, ** P value < 0.01. Mann-Whitney test for **A**.

**Fig S7.**
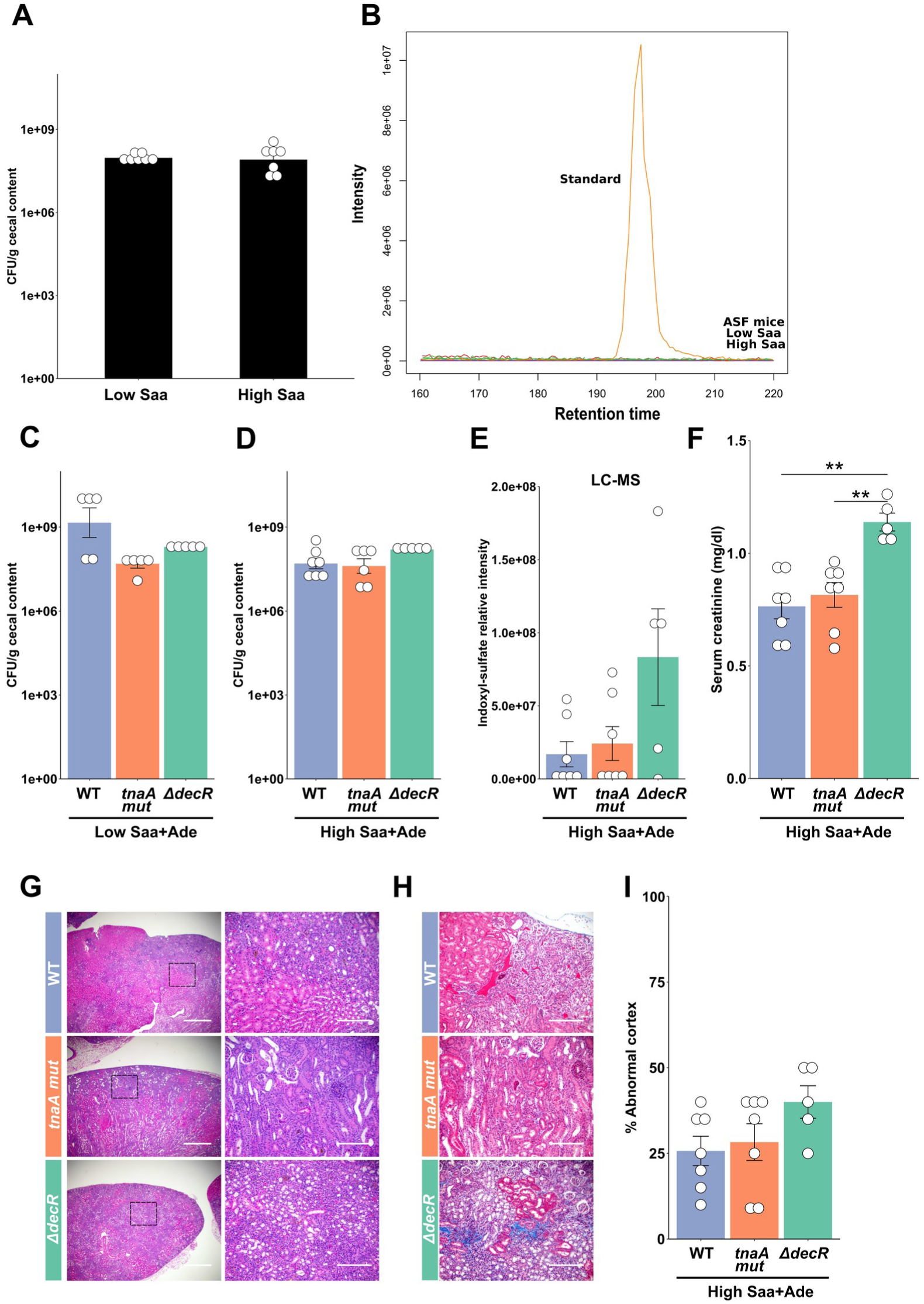
Supplementary data for Fig 4. **A.** Colonization of ASF mice with WT *E. coli* on Saa diets. **B.** Chromatogram of indole detection in cecal contents from ASF mice on Saa diets. **C-D.** Colonization of ASF mice with different *E. coli* strains on low (**C**) and high (**D**) Saa+Ade diet. **E.** LC-MS measurements of serum indoxyl sulfate in ASF mice on high Saa+Ade diet, colonized with *E. coli* strains. **F.** Serum creatinine levels of mice in E. **G.** Representative H&E staining of kidneys from mice in E. **H.** Representative trichrome staining of kidneys from mice in E. **I.** Histology-based renal injury score of mice in E. Data represent 3 independent experiments for **A**, **B**, **C**, **D**, **E**, **F** and **I**. Symbols represent individual mice. Bars represent mean ± SEM. ** P value < 0.01 Two-way ANOVA with Tukey’s post-hoc test for **F.**

**Fig S8.**
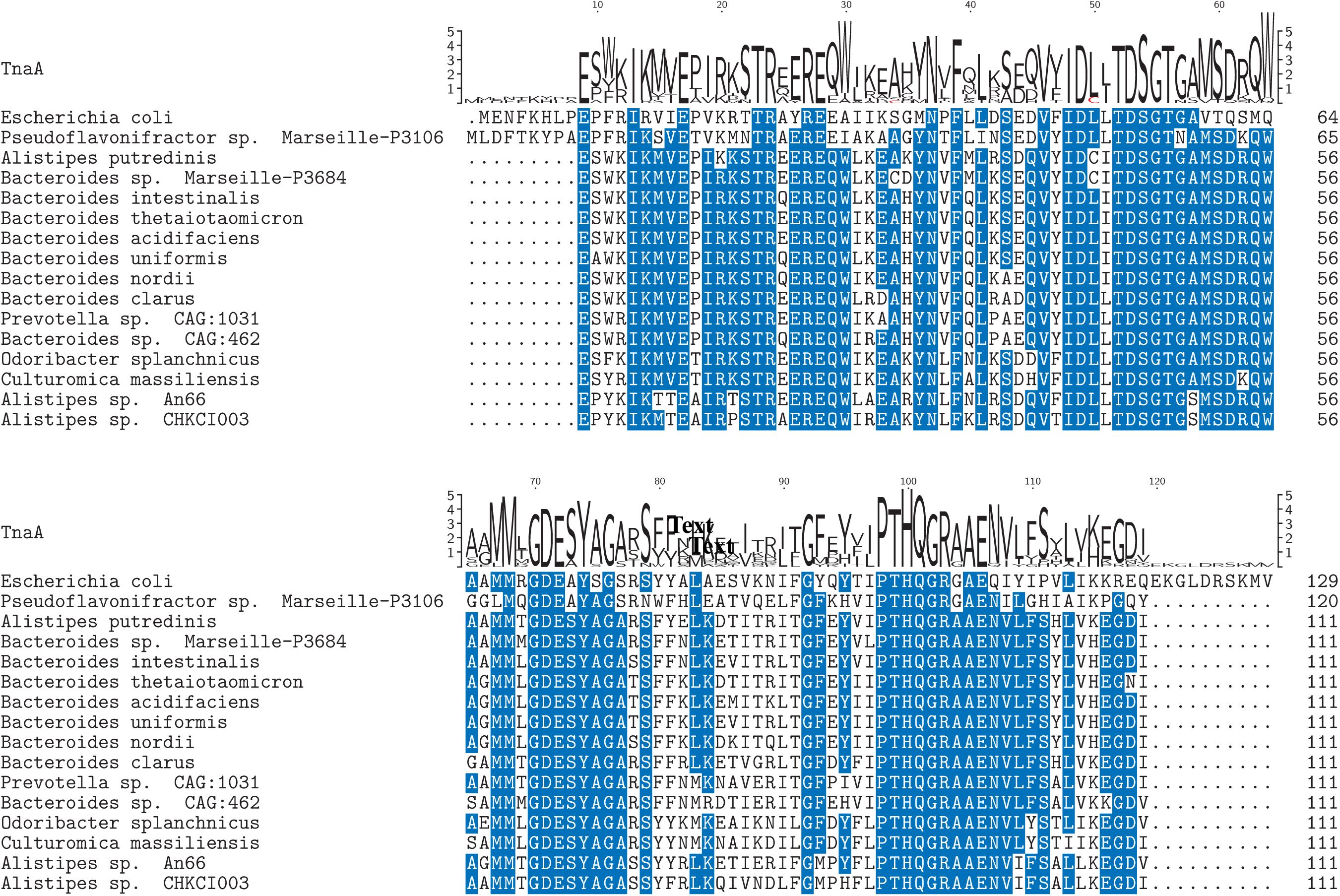
Multiple sequence alignment of TnaA from several human gut microbiota bacterial species. A multiple sequence alignment (MSA) of the 15 closest orthologs of E. coli TnaA in gut bacteria species, obtained by BLAST search with E. coli TnaA vs the microbiome reference sequence database. Identical amino acids are highlighted in blue. The top logo sequence represents the consensus sequence of the MSA with the cysteine residues labeled in red. The MSA was performed using the Clustal Omega algorithm.

